# Depletion of TDP-43 exacerbates tauopathy-dependent brain atrophy by sensitizing vulnerable neurons to caspase 3-mediated endoproteolysis of tau in a mouse model of Multiple Etiology Dementia

**DOI:** 10.1101/2024.06.26.600814

**Authors:** Meghraj S Baghel, Grace D Burns, Margarita Tsapatsis, Aswathy Peethambaran Mallika, Anna Lourdes F Cruz, Tianyu Cao, Xiaoke K Chen, Isabel De La Rosa, Shaelyn R Marx, Yingzhi Ye, Shuying Sun, Tong Li, Philip C Wong

## Abstract

TDP-43 proteinopathy, initially disclosed in amyotrophic lateral sclerosis (ALS) and frontotemporal dementia (FTD), coexists with tauopathy in a variety of neurodegenerative disorders, termed multiple etiology dementias (MEDs), including Alzheimer’s Disease (AD). While such co-pathology of TDP-43 is strongly associated with worsened neurodegeneration and steeper cognitive decline, the pathogenic mechanism underlying the exacerbated neuron loss remains elusive. The loss of TDP-43 splicing repression that occurs in presymptomatic ALS-FTD individuals suggests that such early loss could facilitate the pathological conversion of tau to accelerate neuron loss. Here, we report that the loss of TDP-43 repression of cryptic exons in forebrain neurons (*CaMKII-CreER;Tardbp^f/f^* mice) is necessary to exacerbate tauopathy-dependent brain atrophy by sensitizing vulnerable neurons to caspase 3-dependent cleavage of endogenous tau to promote tauopathy. Corroborating this finding within the human context, we demonstrate that loss of TDP-43 function in iPSC-derived cortical neurons promotes early cryptic exon inclusion and subsequent caspase 3-mediated endoproteolysis of tau. Using a genetic approach to seed tauopathy in *CaMKII-CreER;Tardbp^f/f^*mice by expressing a four-repeat microtubule binding domain of human tau, we show that the amount of tau seed positively correlates with levels of caspase 3-cleaved tau. Importantly, we found that the vulnerability of hippocampal neurons to TDP-43 depletion is dependent on the amount of caspase 3-cleaved tau: from most vulnerable neurons in the CA2/3, followed by those in the dentate gyrus, to the least in CA1. Taken together, our findings strongly support the view that TDP-43 loss-of-function exacerbates tauopathy-dependent brain atrophy by increasing the sensitivity of vulnerable neurons to caspase 3-mediated endoproteolysis of tau, resulting in a greater degree of neurodegeneration in human disorders with co-pathologies of tau and TDP-43. Our work thus discloses novel mechanistic insights and therapeutic targets for human tauopathies harboring co-pathology of TDP-43 and provides a new MED model for testing therapeutic strategies.

**Highlights:**

- Loss of TDP-43 repression of cryptic exons is necessary for caspase 3-dependent endoproteolysis of tau at D421 in the mouse brain and human iPSC-derived cortical neurons.
- The level of caspase 3-dependent cleavage of tau is a major determinant of the vulnerability of mouse brain neurons lacking TDP-43.
- In a novel mouse model of multiple etiology dementia, TDP-43 loss-of-function exacerbates tauopathy-dependent brain atrophy by sensitizing vulnerable neurons to caspase 3-mediated endoproteolysis of tau to drive tauopathy.
- In human tauopathies with co-pathology of TDP-43, dysfunction of TDP-43 may promote caspase 3-dependent cleavage of endogenous tau in vulnerable neurons and exacerbate tauopathy-dependent neurodegeneration.

**Summary:** The pathogenic mechanism by which TDP-43 loss of repression function exacerbates tauopathy-dependent neurodegeneration in multiple etiology dementia (MED) with co-pathology of TDP-43 is unknown. In a novel mouse model of MED, loss of TDP-43 function exacerbates tauopathy-dependent brain atrophy by sensitizing vulnerable neurons to caspase 3-dependent cleavage of endogenous tau to drive tauopathy. This mechanistic insight informs novel targets and therapeutic strategies for MEDs harboring the co-pathologies of tau and TDP-43, which can be validated using this mouse model of MED.

## Introduction

Alzheimer’s disease (AD), the most common age-related dementia and 5^th^ most prevalent cause of death in individuals over 65 in the United States, is associated with progressive loss of synapses, and eventually neurons (Terry et al., 1991). While strong evidence supports a linear disease progression triggered by excessive amyloid-β (Aβ) in familial AD, multifactorial etiology has been hypothesized for sporadic late onset AD (Self et al., 2023; De Strooper et al., 2016; Karran et al., 2022; Long et al., 2019; Martens et al., 2022; Zhou et al., 2020; Chen et al., 2022; Sun et al., 2023; Mathys et al., 2019; Blanchard et al., 2022; Martorell et al., 2019; Mathys et al., 2023). Recent studies have demonstrated that up to 75% of AD and AD-Related Dementia (ADRD) cases exhibit non-canonical pathologies, including TAR DNA/RNA binding protein (TDP-43), α-synuclein “Lewy bodies” and tau “pick bodies” (Josephs et al., 2014; Josephs et al., 2015; Robinson et al., 2018; Neumann et al., 2006; Probst al., 1996; Stefanis, 2012). TDP-43 pathology is observed in 40-60% of ADRD cases and is strongly associated with worsened neurodegeneration and cognition (Josephs et al., 2014; Josephs et al., 2015; Robinson et al., 2018). TDP-43 pathology is also observed in some cases of tauopathies including frontotemporal lobar dementia (FTLD) (Kim et al., 2018), corticobasal degeneration (CBD) (Koga et al., 2018), primary age-related tauopathy (PART) (Josephs et al., 2017) and progressive supranuclear palsy (PSP) (Yokota et al., 2010). Therefore, it is likely that the occurrence and interaction of these pathological factors lead to worsened neurodegeneration and steeper cognitive decline.

TDP-43 is centrally associated with ALS-FTD (Neumann et al., 2006) and regulates the repression of non-conserved cryptic exons (Ling et al., 2015). Genetic, fluid biomarker and neuropathological findings support the notion that loss of TDP-43 splicing repression begins presymptomatically to drive neuron loss in ALS-FTD and AD-TDP (Vatsavayai et al., 2016; Sun et al., 2017; Irwin et al., 2023; Seddighi et al., 2023; Melamed et al., 2019; Klim et al., 2019; Brown et al., 2022; Ma et al., 2022). While the clinical significance of TDP-43 pathophysiology is undisputed, evidence for pathogenic mechanisms as to how loss of TDP-43 accelerates neuron loss in MEDs is currently lacking.

Studies have reported neuronal co-localization and interaction of phosphorylated tau (p-tau) and TDP-43 positive inclusions in AD brains (Tomé et al, 2021), but the exact nature of co-localization remains elusive. Co-expression of tau and TDP-43 in animal models was shown to promote tau phosphorylation, suggesting synergism between tau and TDP-43 (Davis et al., 2017; Latimer et al., 2022). Recent findings using a human autopsy cohort, tau biosensor line, and mutant TDP-43 mouse model support the idea that TDP-43 pathology correlates with more severe tauopathy (Tomé et al., 2023).

We and others previously demonstrated that Aß plaque deposition is one necessary factor that facilitates the pathological conversion of endogenous tau stimulated by a human four-repeat domain (hTauRD) tau seed to drive neuron loss in an age-dependent manner (Li et al., 2016), which also occurs in the human AD brain. Likewise, intracerebral injection of AD-tau seeds facilitated Aß plaque-dependent misfolding of endogenous tau (He et al., 2018). Moreover, caspase 3-dependent cleavage of tau in the AD brain and mouse models has been well-documented (Gervais et al., 1999; Rohn et al., 2001; Su et al., 2001; Gastard et al., 2003; Guo et al., 2004; Calignon et al., 2010).

Based on these findings, coupled with our previous observations that loss of TDP-43 in forebrain neurons (*CaMKII-CreER;Tardbp^f/f^* mice) leads to selective death of CA2/3 neurons accompanied by modest activation of caspase 3 (LaClair et al, 2016), we hypothesize that loss of TDP-43 function is a key determinant necessary to sensitize vulnerable neurons to caspase 3-dependent cleavage of tau and accelerate tauopathy-dependent neurodegeneration in MEDs harboring TDP-43 pathology. To test these notions, we first established that inclusion of TDP-43 cryptic exons leads to activation of caspase 3-mediated endoproteolysis of tau in both mouse and human model systems. Subsequently, we took a genetic approach to elevate the level of caspase 3-dependent cleavage of tau in our TDP-43 conditional knockout mouse model (*CaMKII-CreER;Tardbp^f/f^*mice) (LaClair et al., 2016) to mimic the loss of TDP-43 function that occurs in MEDs with co-pathology of TDP-43. We created *CaMKII-CreER;Tardbp^f/f^* mice expressing a tau seed (Li et al., 2016) in which a human tau fragment encompassing the four-repeat domain of tau (*hTauRD*) is expressed in central neurons to levels similar to that of endogenous tau (*CaMKII-CreER;Tardbp^f^*^/*f*^*;hTau4R* mice). To achieve an even higher level of caspase 3-dependent cleavage of tau, we intraparenchymally delivered an adeno-associated viral vector expressing the *hTauRD* transgene (AAV-PhP.eB-hTauRD) to prime tauopathy in the hippocampus of *CaMKII-CreER;Tardbp^f^*^/*f*^ mice. Using these two complementary approaches, we elucidate the pathogenic mechanism whereby loss of TDP-43 function promotes tauopathy and death of vulnerable neurons that are sensitive to levels of caspase 3-dependent cleavage of tau.

## Results

### Inclusion of TDP-43 dependent cryptic exons precedes caspase 3-mediated cleavage of tau and leads to selective death of hippocampal excitatory neurons

Previously, we showed that the deletion of TDP-43 in forebrain excitatory neurons led to selective loss of neurons in hippocampal CA2/3 accompanied by modest activation of caspase 3 (LaClair et al, 2016). To determine whether loss of TDP-43 splicing repression precedes caspase 3 activation, we first confirmed that eight months after depletion of TDP-43 within the hippocampal circuit, as compared to littermate controls, CA2/3 neurons were most vulnerable to the loss of TDP-43 in aged (19-month-old) *CaMKII-CreER;Tardbp^f/f^* mice (Fig. 1A). We observed selectively in neurons of CA2/3, but not those in CA1 or DG, that the activation of caspase 3 occurs one month after the depletion of TDP-43 (Fig. 1B); subsequently, caspase 3 is activated in CA1 and DG neurons two months after depletion of TDP-43 (Fig. 1B, third column; Sup Fig. 2A-B) and accumulated over time (Fig. 1B, right column; Sup Fig. 2A, middle column). Furthermore, after neuron loss occurred in CA2/3, the level of caspase 3 returned to baseline (Fig. 1B, right column;Sup Fig. 2A, middle column). To directly test whether inclusion of cryptic exons (Ling et al., 2015) precedes activation of caspase 3, we assessed inclusion of a TDP-43 dependent cryptic exon within the mouse *Unc13a* pre-mRNA (Jeong et al., 2017). One month after depletion of TDP-43, we observed robust inclusion of this cryptic exon not only in CA2/3 neurons of *CaMKII-CreER;Tardbp^f/f^* mice, but also those within CA1 and DG as assessed by a BaseScope probe that specifically identifies the *Unc13a* cryptic exon (Fig. 1C, middle column). These findings strongly support the view that inclusion of cryptic exons precedes the activation of caspase 3 in *CaMKII-CreER;Tardbp^f/f^*mice lacking TDP-43 in excitatory hippocampal neurons.

**Figure 1.**
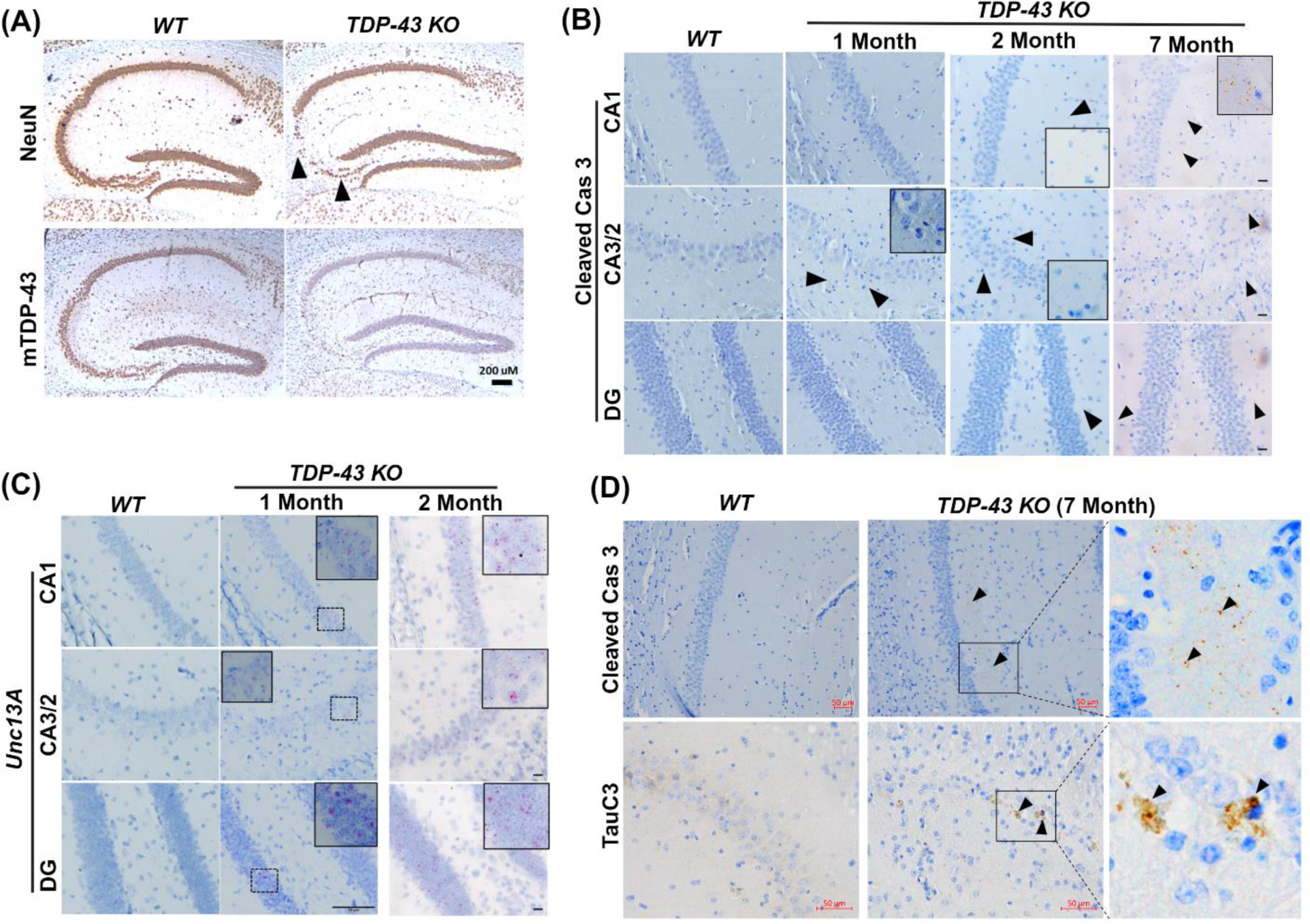
Loss of TDP-43 repression function leads to selective vulnerability for neuron loss in hippocampal CA2/3 associated with caspase 3-mediated cleavage of tau. **(A)** Immunohistochemical analysis of hippocampus in sagittal sections of brains of 19-month-old *WT* (n=3), and *TDP-43 KO* (n=6) mice. The upper panel shows immunohistochemical staining using an antibody specific to NeuN to detect neurons and the lower panel shows images using an antibody specific to mouse TDP-43 revealing depletion of TDP-43 in the mouse hippocampus. Arrow heads indicated selective loss of neurons in CA2/3 (scale bar, 200μm) **(B)** Cleaved caspase 3 immunohistochemistry in brain sections of TDP-43 deleted and WT mice at different time points (1, 2 and 7 months) after TDP-43 deletion. Enlarged inset view of immunoreactive area, arrow heads show the positive signal in CA1, CA2/3 and DG subregions of hippocampus (scale bar, 20µm). **(C)** BaseScope analysis of *Unc13a* cryptic exon in CA2/3, CA1 and DG subregions of hippocampus in WT and TDP-43 depleted mice. (Left and middle column, Scale bar 50µm; right column scale bar, 20µm). **(D)** Immunohistochemical analysis of *WT* (n=3), and *TDP-43 KO* (n=6) mice depleted of TDP-43 at 12 months of age and analyzed at 19-months-old. Upper panel shows cleaved caspase 3 immunoreactivity in CA1 subregion of hippocampus, arrow heads indicate positive signal in middle and enlarged right column of boxed area (Scale bar, 50µm). Lower panel depicts the immunoreactivity of caspase cleaved tau (TauC3), arrow heads show positive signal in middle and enlarged right column of boxed area (Scale bar, 50µm)

**Figure 2.**
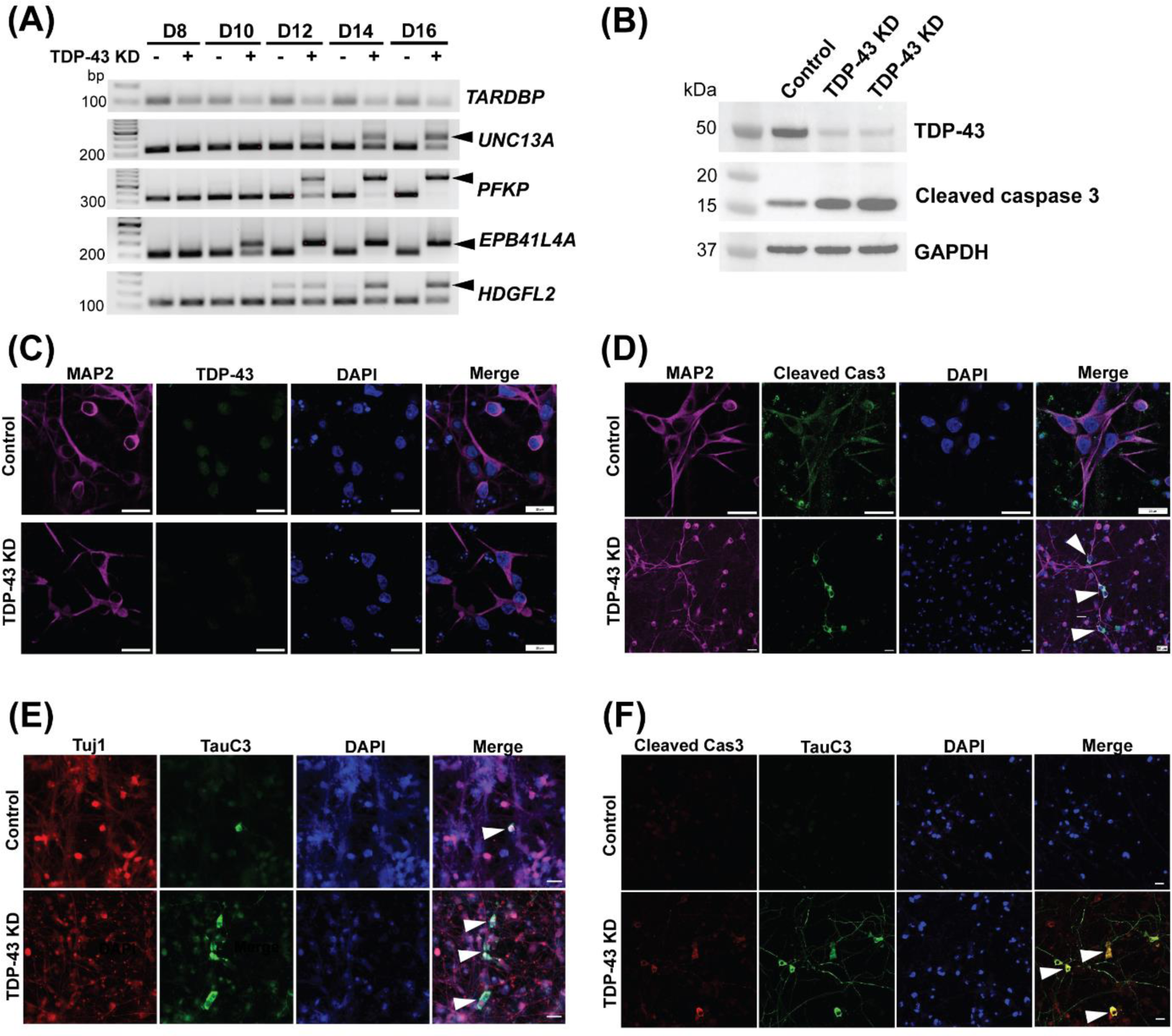
Inclusion of cryptic exon precedes caspase 3 activation leading to cleavage of tau in human neurons deficient in TDP-43. **(A)** RT-PCR analysis of TDP-43 cryptic exon targets from day 8, 10, 12, 14 and 16 i3Neurons showing cryptic exon expression begins as early as day 10. **(B)** Immunoblot showing efficient knockdown of TDP-43 and cleaved caspase 3 expression at day 15 in i3Neurons treated with a scrambled RNA lentivirus or lentivirus targeting TDP-43. **(C)** Representative immunofluorescence images of 14-day-old i3Neurons using antibodies against MAP2 and TDP-43 (scale bar, 20µm). **(D)** Representative immunofluorescence images of 14-day-old i3Neurons using antibodies against MAP2 and cleaved caspase 3 to show activated caspase 3 in TDP-43 depleted cells (scale bar, 20µm). **(E)** Representative immunofluorescence images of 14-day-old i3Neurons using antibodies against Tuj1 and caspase-3-cleaved tau (TauC3) showing increased cleaved tau in TDP-43 depleted cells (scale bar, 20µm). **(F)** Co-immunofluorescence of cleaved caspase 3 and TauC3 showing colocalization in ∼50% neurons (scale bar, 20µm).

Given that caspase activation dependent cleavage of tau precedes tau tangle formation (Calignon et al., 2010), we asked whether cleavage of endogenous tau occurs in CA2/3 neurons of *CaMKII-CreER;Tardbp^f/f^* mice during aging. Importantly, loss of TDP-43 function led to the activation of caspase 3 (cleaved caspase 3) dependent cleavage of endogenous tau (using antisera, TauC3, recognizing the neoepitope exposed by caspase 3 at position D421 of tau) (Fig. 1D).

### Inclusion of cryptic exons in human cortical neurons deficient in TDP-43 precedes caspase 3 –mediated cleavage of tau

To corroborate our finding that loss of TDP-43 repression of cryptic exons occurs prior to caspase 3 activation in human neurons, we depleted TDP-43 through exposure of lentivirus expressing an shRNA targeting TDP-43 (shTDP-43) in iPSC-derived i3Neurons (Tian et al., 2019). As assessed by RT-PCR, immunoblot and immunofluorescence analysis, we observed efficient knockdown of TDP-43 as compared to the non-targeting, scrambled control (Fig. 2A-C). Intriguingly, inclusion of various cryptic exons was detected as early as day 10 upon TDP-43 depletion (Fig. 2A). By day 15, we observed a baseline level of caspase 3 activation in control i3Neurons but marked activation in TDP-43 depleted neurons (cleaved caspase 3, 17kDa; Fig. 2B & D). These findings support the notion that cryptic exon inclusion precedes caspase activation. As expected, we showed that loss of TDP-43 induces increased cleavage of endogenous tau as observed by TauC3 immunofluorescence staining (Fig. 2E). Since caspase 3 activation precedes tau cleavage, we confirmed the colocalization of cleaved caspase 3 and C-terminal truncated tau (Fig. 2F). Together, these findings support the notion that inclusion of TDP-43 dependent cryptic exons promotes caspase 3 mediated endoproteolysis of tau, which primes tau misfolding and propagation, leading to loss of vulnerable neurons in human disease exhibiting the co-pathology of tau and TDP-43.

### TDP-43 loss-of-function facilitates age-dependent death of dentate granule neurons in the presence of a tau seed

That only CA2/3 neurons are vulnerable to the loss of TDP-43 within the hippocampal circuit in *CaMKII-CreER;Tardbp^f/f^* mice raised an interesting question as to whether other co-pathologic factors are required to activate caspase 3 above a certain threshold that determines the vulnerability of other excitatory neurons. Previously, we showed that hTauRD is insufficient to drive the pathological tau conversion in mice, but it could seed the conversion of endogenous tau to drive tauopathy and neuron loss in an age-dependent manner in the presence of Aβ plaques (Li et al., 2016). We reasoned that such a tau seeding model would be another trigger for revealing other vulnerable neurons within the hippocampal circuit when TDP-43 is depleted, mimicking a human disease context for this type of MED with co-pathologies of tau and TDP-43. To determine the potential interactions of TDP-43 and tau pathologies, a crossbreeding strategy with *CaMKII-CreER;Tardbp^f/f^* (TDP-43 KO) and *hTau4R* mice was employed to generate a cohort of *CaMKII-CreER;Tardbp^f/f^*;*hTau4R* mice along with a set of littermate controls (Sup Fig. 1). We deleted *Tardbp* selectively in excitatory forebrain neurons at 12 months-of-age and analyzed these mice at various times up to 25 months (Fig. 1A). As expected, brain extracts from these mice subjected to immunoblot analysis confirmed the accumulation of the hTau4RD fragment (∼16kDa) specifically expressed in *Tau4R* and *CaMKII-CreER;Tardbp^f/f^*;*hTau4R*, but not control, mice (Fig. 3A).

**Figure 3.**
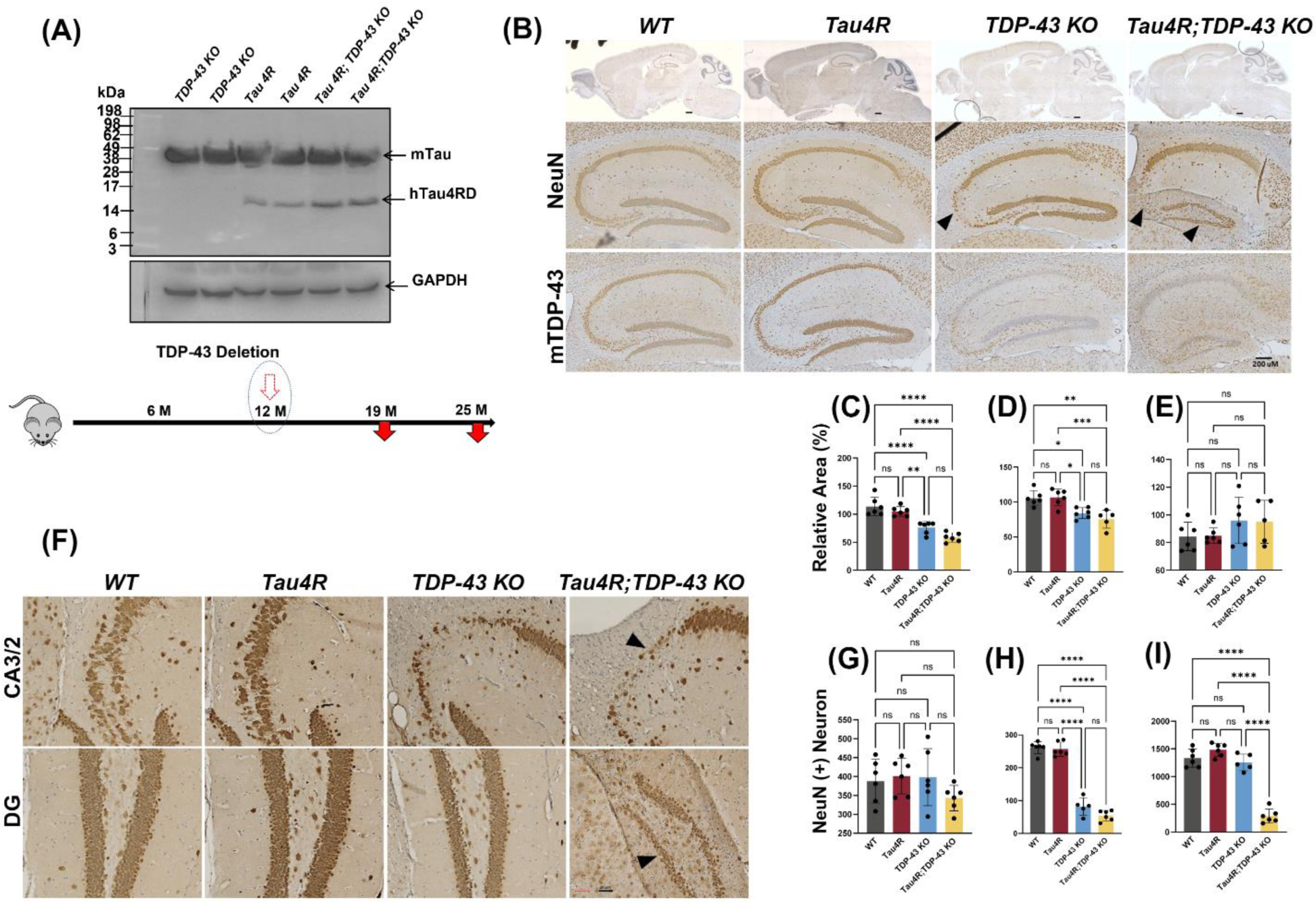
Loss of TDP-43 function accelerates neurodegeneration in hTau4R expressing mice. **(A)** Immunoblot using a 77G7 antibody which recognizes the specific expression of hTau4RD fragments (∼16kDa) including endogenous mouse tau protein expression. Bottom diagram shows the timeline for induction whereby TDP-43 is deleted at 12 months of age and mice are analyzed at either 19 or 25 months of age. **(B & F)** Immunohistochemical analysis of brains of 19-month-old *WT* (n=6), *Tau4R* (n=6), *TDP-43 KO* (n=5) and *Tau4R;TDP-43 KO* mice (n=6) using antiserum specific to NeuN to detect neurons; sagittal sections of whole brains (upper panel of B; scale bar, 500μm) and hippocampi (lower panels of B; scale bar, 200μm), F; magnified views of NeuN immunohistochemistry in hippocampal subregions CA2/3 & DG arrow heads indicates loss of neurons (both panels; scale bar, 20μm). Lower panel of B; immunohistochemistry of mouse TDP-43 showing depleted TDP-43 in mouse hippocampus (scale bar, 200μm). **(C-E)** Analysis of relative area measurements of hippocampus, cortex and cerebellum respectively depicted (% relative area of *WT*) (one-way ANOVA; ns: no significant difference; *P=0.0111 (*WT vs TDP-43 KO*) *P=0.0168 (*Tau4R vs TDP-43 KO*) **P=0.0014; ****P<0.0001). (**G-I**) Neuronal cell count of CA1, CA2/3 and DG regions from 19-month-old cohort (one-way ANOVA; ns: no significant difference; ****P<0.0001).

While depletion of TDP-43 led to selective loss of neurons in the CA2/3 subregion of the hippocampus of *CaMKII-CreER;Tardbp^f^*^/*f*^ mice as previously seen (LaClair et al., 2016), we observed neuron loss extending beyond this region in *CaMKII-CreER;Tardbp^f^*^/*f*^*;hTau4R* mice (Fig. 3B, F). As compared to age-matched littermates, *CaMKII-CreER;Tardbp^f^*^/*f*^*;hTau4R* mice also exhibited selective loss of granule neurons in the DG (Fig. 3F). To analyze neuronal viability, we quantified the number of NeuN positive neurons in hippocampal subregions CA1, CA3/2 and DG and showed that while loss of TDP-43 in *CaMKII-CreER;Tardbp^f^*^/*f*^ mice promoted selective loss of neurons in CA2/3 (Fig. 3H) as expected, presence of the tau seed additionally triggered neuron loss in DG of *CaMKII-CreER;Tardbp^f^*^/*f*^*;hTau4R*, but not in age-matched control mice (Fig. 3I). However, there is no quantitative difference found in CA1 neuronal populations (Fig. 3G). Moreover, we found that while the relative areas of hippocampus and cortex were significantly decreased in *CaMKII-CreER;Tardbp^f^*^/*f*^ mice (as compared to hTau4R transgenic and wild-type littermates), this reduction was further exacerbated in *CaMKII-CreER;Tardbp^f^*^/*f*^*;hTau4R* mice (Fig. 3C-D). As expected, there was no significant difference in the relative area of the cerebellum amongst all genotypes (Fig. 3E). These results are consistent with the idea that loss of TDP-43 function which reflects nuclear clearance of TDP-43 occurring in human disease, promotes the exacerbated neurodegeneration occurring in MED subtypes exhibiting co-pathologies of TDP-43 and tau.

### Vulnerability of dentate granule neurons to TDP-43 depletion is sensitive to caspase 3-dependent cleavage of tau

Loss of dentate neurons due to TDP-43 depletion in *CaMKII-CreER;Tardbp^f^*^/*f*^*;hTau4R* mice suggests the possibility that increased level of caspase 3-dependent cleavage of tau facilitated by hTauRD underlies neuronal vulnerability within the hippocampal circuit. To test this notion, we assessed caspase 3 activation in our model of MED exhibiting TDP-43 and tau pathologies. Since caspase 3 activation resulting in tau cleavage and aggregation has been linked to AD (Gervais et al, 1999; Rohn et al 2001; Su et al 2001; Gastard et al 2003; Guo et al 2004; Rissman et al, 2004), we questioned whether this phenomenon was at play in our MED model. That depletion of TDP-43 in the presence of a tau seed sensitized dentate granule neurons to cell death led us to examine whether caspase 3 activation is elevated. As compared to *CaMKII*-*CreER;Tardbp^f^*^/*f*^ mice, a marked increase of activated caspase 3 was observed in dentate neurons of *CaMKII-CreER;Tardbp^f^*^/*f*^*;hTau4R* mice (Fig. 4A), suggesting that the vulnerability of hippocampal neurons depends on the level of activated caspase 3. Indeed, while activation of caspase 3 in the hippocampus occurred in *CaMKII*-*CreER;Tardbp^f^*^/*f*^ mice between 19 and 25 months-of-age, such caspase 3 activation was markedly accelerated in *CaMKII-CreER;Tardbp^f^*^/*f*^*;hTau4R* mice (Fig. 4B); this effect was age dependent as 25-month-old mice exhibited enhanced caspase 3 activation as compared to 19-month-old mice (Fig. 4B).

**Figure 4.**
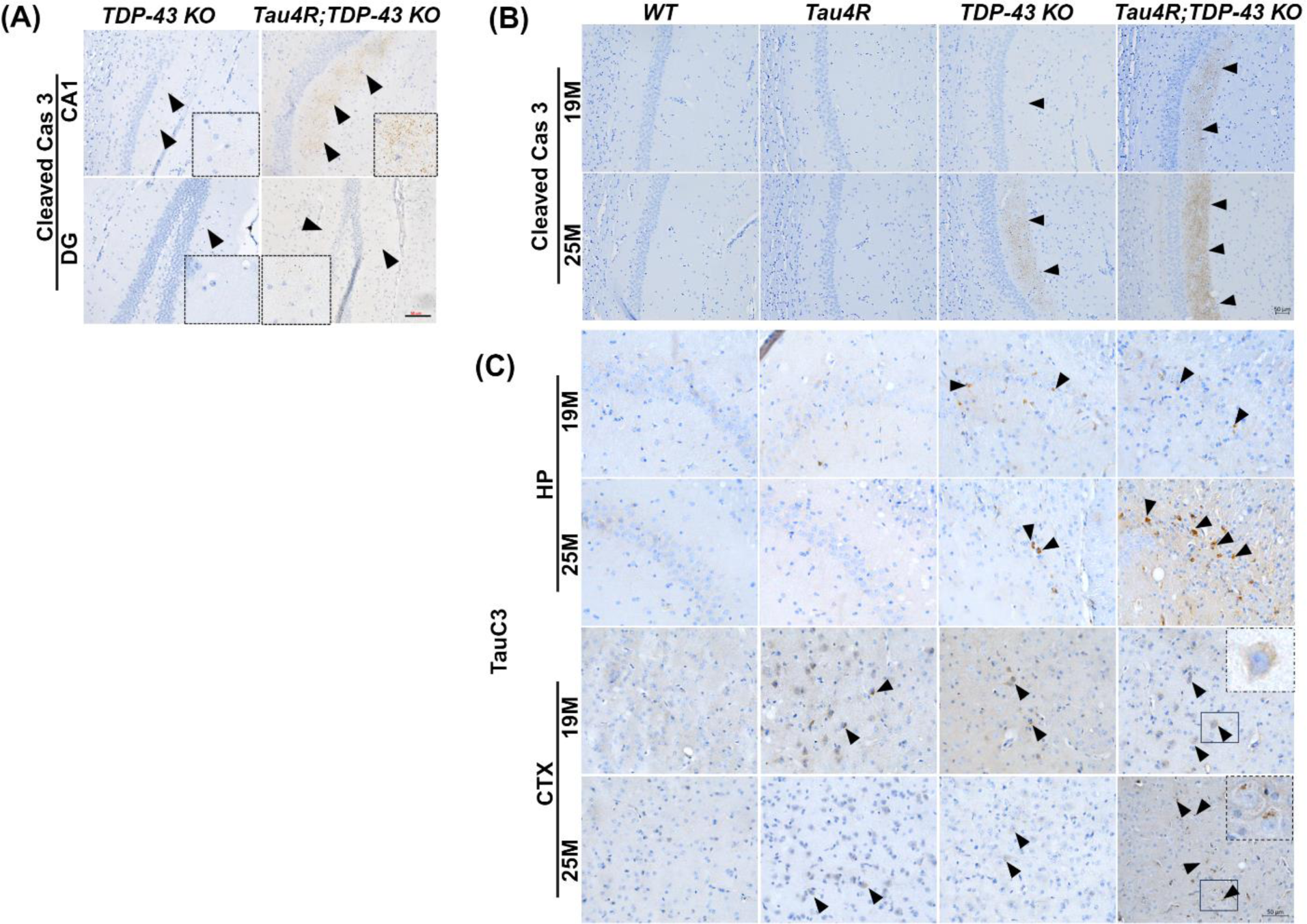
Caspase 3 is activated leading to cleavage of tau in hTau4R-expressing mice lacking TDP-43. **(A)** Immunohistochemical analysis of cleaved caspase 3 in brain sections of 16-month-old old mice deleted at 12 months of age *TDP-43* KO (n=3) and *Tau4R;TDP-43 KO (*n=3) mice; inset enlarged image shows positive staining (scale bar, 50μm). (**B**) Immunohistochemical analysis of brain sections of 19-month-old *WT* (n=6), *Tau4R* (n=6), *TDP-43 KO* (n=5) and *Tau4R;TDP-43 KO* mice (n=6) or 25-month-old *WT* (n=3), *Tau4R* (n=4), *TDP-43 KO* (n=5) and *Tau4R;TDP-43 KO* mice (n=4) in the CA1 region of the hippocampus using antiserum against cleaved caspase 3 (scale bar, 50µm) **(C)** caspase-cleaved tau (TauC3) in the CA2/3 region of the hippocampus (HP) (CA2/3) and cortex (CTX) (scale bar, 50μm).

To determine whether the caspase-mediated cleavage of tau is elevated in hippocampal neurons of *CaMKII-CreER;Tardbp^f^*^/*f*^*;hTau4R* mice we employed an antibody recognizing a neo-epitope of tau-D421. Immunohistochemical analysis revealed the presence of cleaved tau in an age dependent manner in hippocampal CA1, CA2/3, and cortex of *hTau4R*, *CaMKII-CreER;Tardbp^f^*^/*f*^ and *CaMKII-CreER;Tardbp^f^*^/*f*^*;hTau4R* mice, but not in non-transgenic controls (Fig. 4C, Sup Fig. 3). As compared to *hTau4R* or *CaMKII-CreER;Tardbp^f^*^/*f*^ mice, a marked elevation of cleaved tau is found in *CaMKII-CreER;Tardbp^f^*^/*f*^ and *CaMKII-CreER;Tardbp^f^*^/*f*^*;hTau4R* mice (Fig. 4C). We additionally confirmed the antibody specificity of cleaved tau at D421 (TauC3) in *CaMKII-CreER;Tardbp^f^*^/*f*^*;hTau4R* mice (Sup Fig. 4, left two columns: negative primary and secondary antibody staining; right column; regular staining of TauC3). These results strongly support the idea that the vulnerability of hippocampal neurons is highly sensitive to the loss of TDP-43 dependent activation of caspase 3 to cleave tau, thus providing a pathogenic mechanism underlying MED harboring co-pathologies of tau and TDP-43.

### Loss of TDP-43 function promotes the pathological conversion of endogenous tau to exacerbate tauopathy-dependent death of vulnerable neurons

That elevated caspase 3 activation and tau cleavage occurs in an age dependent manner in *CreER;Tardbp^f^*^/*f*^*;hTau4R* mice and loss of TDP-43 exacerbates neurodegeneration in the hippocampus of 25-month-old *CaMKII-CreER;Tardbp^f^*^/*f*^*;hTau4R* mice (Fig. 5A) raised the question as to whether TDP-43 loss can facilitate pathological conversion of endogenous tau. Immunohistochemical analysis of aged (25-month-old) *CaMKII-CreER;Tardbp^f^*^/*f*^*;hTau4R* mice using two independent antisera (AT8 & pS422) to recognize hyperphosphorylated pathological tau tangles derived from endogenous mouse tau revealed that loss of TDP-43 facilitates age-dependent pathological conversion of wild-type tau in hippocampus (HP), cortex (CTX) and particularly within the entorhinal cortex (EC) (Fig. 5B-C, Sup Fig. 5) where tauopathy is thought to initiate in brains of cases of AD. Importantly, we observed pathological conversion of tau in these mice only after 19 months-of-age (Sup Fig. 6) accompanied by an age-dependent increase of hTauRD (Fig. 5D; Sup Fig. 7). These results support the view that TDP-43 depletion occurring in MED activates caspase 3-mediated cleavage of tau, sensitizes vulnerable neurons to tauopathy and exacerbates tauopathy-dependent neuron loss.

**Figure 5.**
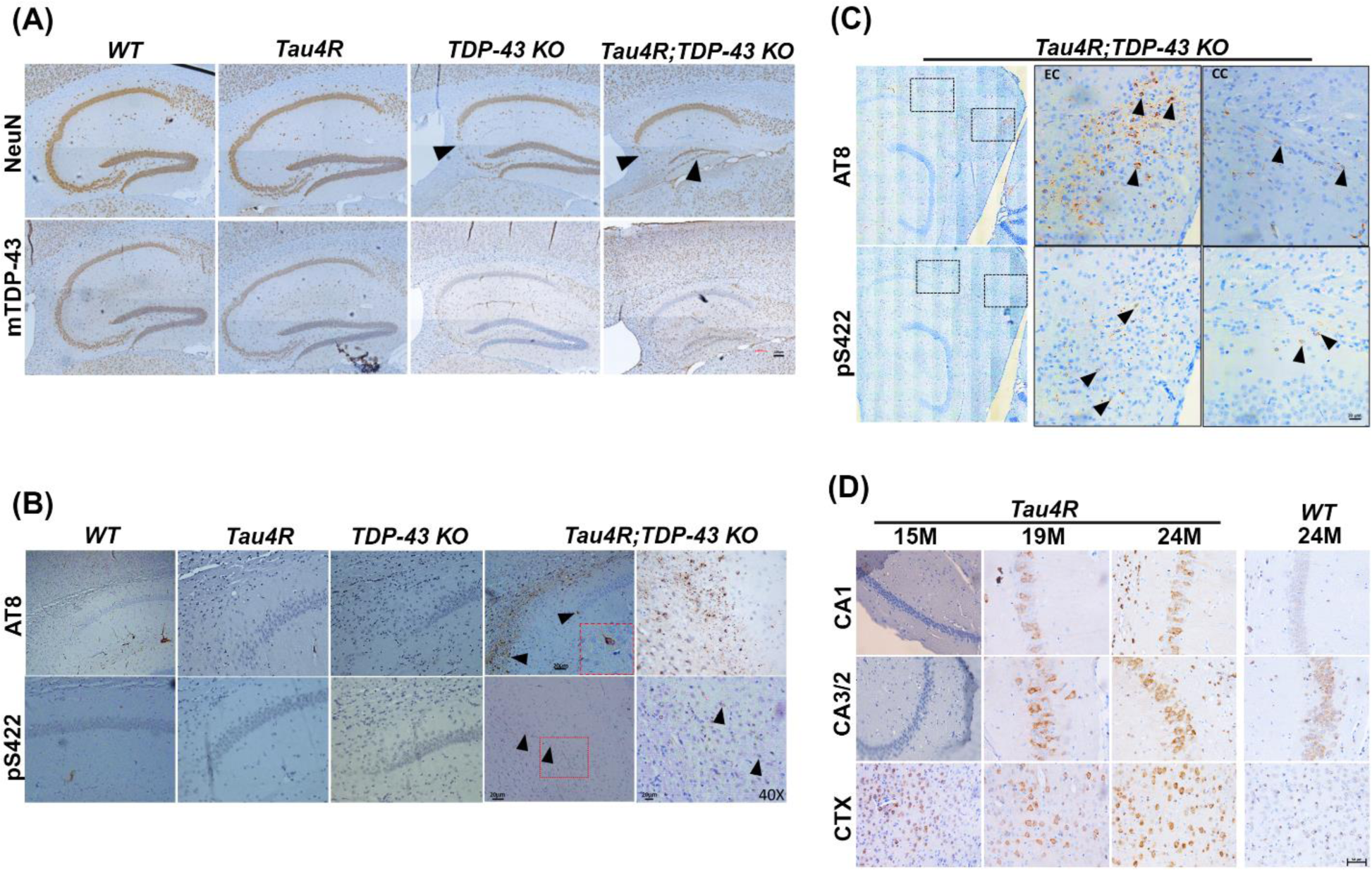
Loss of TDP-43 function facilitates pathological conversion of wild type endogenous tau in hTau4R expressing aged mice. **(A)** Immunohistochemical analysis of the hippocampus in sagittal sections of brains of 25-month-old *WT* (n=3), *Tau4R* (n=4), *TDP-43 KO* (n=5) and *Tau4R;TDP-43 KO* mice (n=4). Upper panel shows immunohistochemical staining using an antibody specific to NeuN to detect neurons and lower panel shows images using an antibody specific to mouse TDP-43 showing depleted TDP-43 in mouse hippocampus (scale bar, 100μm). (**B**) Immunohistochemical analysis of the hippocampus using antibodies specific to endogenous phosphorylated tau, AT8 (upper panel) and pS422 (lower panel). Arrow heads show pathological conversion in *Tau4R;TDP-43 KO* mice (scale bar, 20μm for 20X followed by magnified view at 40X). The sections were counterstained with hematoxylin (blue). (**C**) To see the first occurrence in entorhinal cortex (EC) representative stitched image showing relative strong immunoreactivity with pathological tau (AT8 & pS422) in *Tau4R;TDP-43* mice (scale bar, 20μm). **(D)** Immunohistochemistry analysis showed an age-dependent increase in the accumulation of hTau4R fragment using 4R repeat recognizing antibody (Tau4R;E7T4F) in subregions of HP and CTX of *Tau4R* mice (aged to 15, 19 & 24 months of age), but not in 24-month-old *WT* mice (right panel). Scale bar, 50 μm.

**Figure 6.**
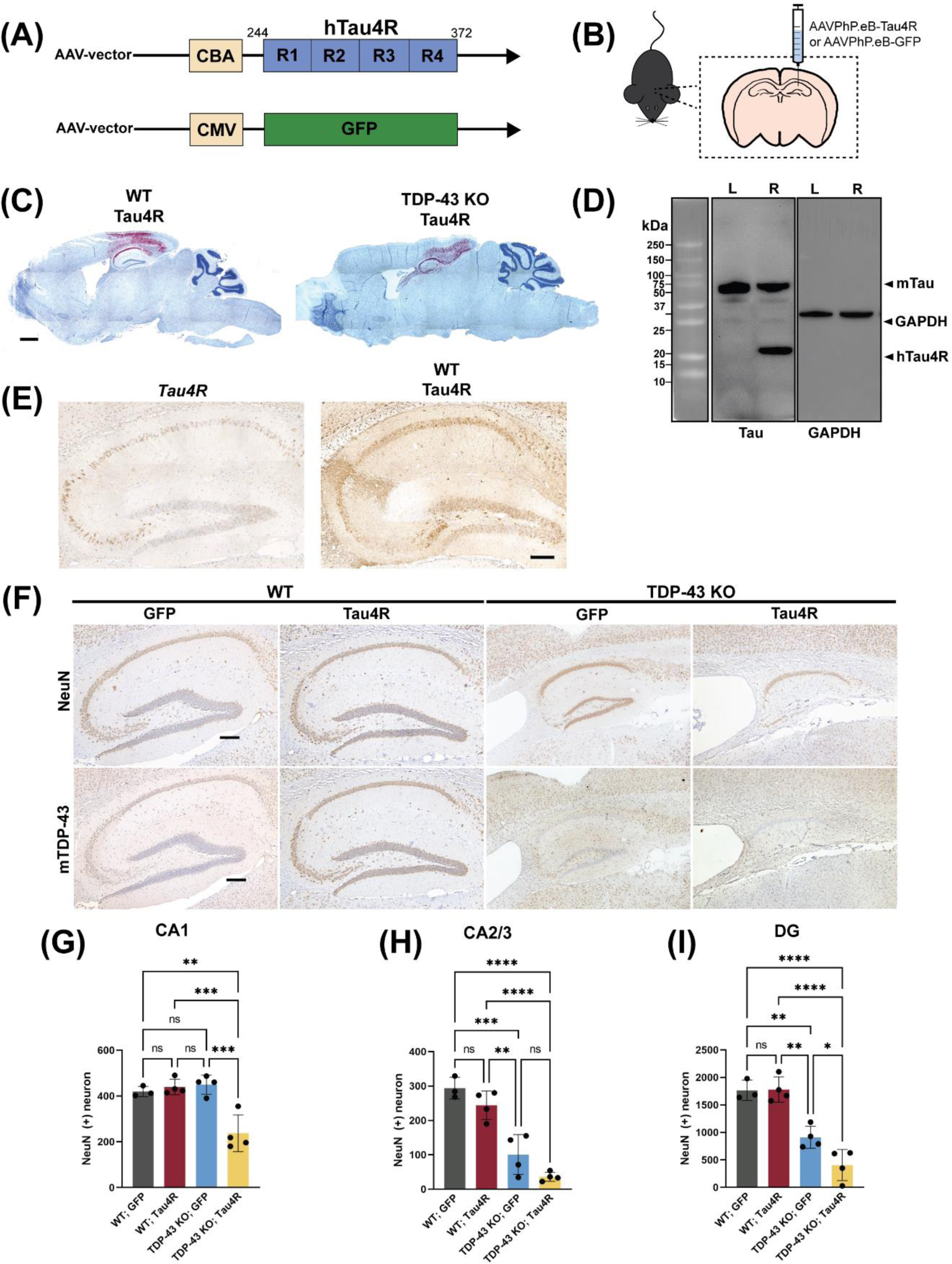
AAV delivery of hTauRD in brains of TDP-43-depleted mice exacerbates neurodegeneration. **(A)** Graphical illustration of the structure of expression vectors packaged into AAV-PhP.eB. The upper schematic diagram shows the construct with the four-repeat domain of human tau (hTau4R) downstream of the chicken β-actin (CBA) promoter. The lower one shows the control vector containing fragment encoding GFP downstream of the cytomegalovirus (CMV) promoter. **(B)** Schematic diagram showing the injection of AAV-PhP.eB-hTau4R or GFP into the right hippocampus. **(C)** BaseScope analysis of whole brain sections from wild-type or TDP-43 KO mice 2 months post injection showing the presence of viral RNA in the hippocampus and region of cortex right above the hippocampus. (scale bar, 1000µm). **(D)** Western blot using 77G7 antibody that recognizes the repeat domain of tau showed the presence of exogenous (∼16kDa) hTau4R and KJ9A antibody for mouse endogenous tau protein from hippocampal lysates of a mouse unilaterally injected in the right hippocampi (R: right; L: left) with hTau4R. Note the expression level of endogenous tau is similar to exogenous in the right hemisphere. **(E)** Representative immunohistochemical analysis of 19-month-old *Tau4R* mice and 12-month-old WT mice sacrificed 6 months post injection with AAV-PhP.eB-hTau4R using an antibody against Tau4R. While several hippocampal cells express Tau4R in *Tau4R* mice, Tau4R expression is greatly increased and more widespread in not only cell bodies but also axons upon AAV delivery (scale bar, 20µm). **(F)** Immunohistochemical analysis of brain sections from 18-month-old WT mice injected with GFP (n=3), WT mice injected with Tau4R (n=4), TDP-43 KO mice injected with GFP (n=4), and TDP-43 KO mice injected with Tau4R (n=4) using antibody against NeuN and mouse TDP-43 (scale bar, 200µm). **(G-I)** Neuronal cell counts of CA1, CA2/3 and DG regions of the hippocampus (one-way ANOVA; ns: no significant difference; CA1 **P=0.0033, ***P=0.0008 (*WT;Tau4R* vs. *TDP-43 KO;Tau4R*), ***P=0.0005 (*TDP-43 KO;GFP* vs. *TDP-43 KO; Tau4R*); CA2/3 **P=0.0018 (*WT;Tau4R* vs. *TDP-43 KO;GFP*), ***P=0.0003 (*WT;GFP* vs. *TDP-43 KO;GFP*), ****P<0.0001; DG *P=0.0440 (*TDP-43 KO;GFP* vs. *TDP-43 KO;Tau4R*), **P= 0.0025 (*WT;GFP* vs. *TDP-43 KO;GFP*), **P=0.0012 (*WT;Tau4R* vs. *TDP-43 KO; GFP*), ****P<0.0001.)

**Figure 7.**
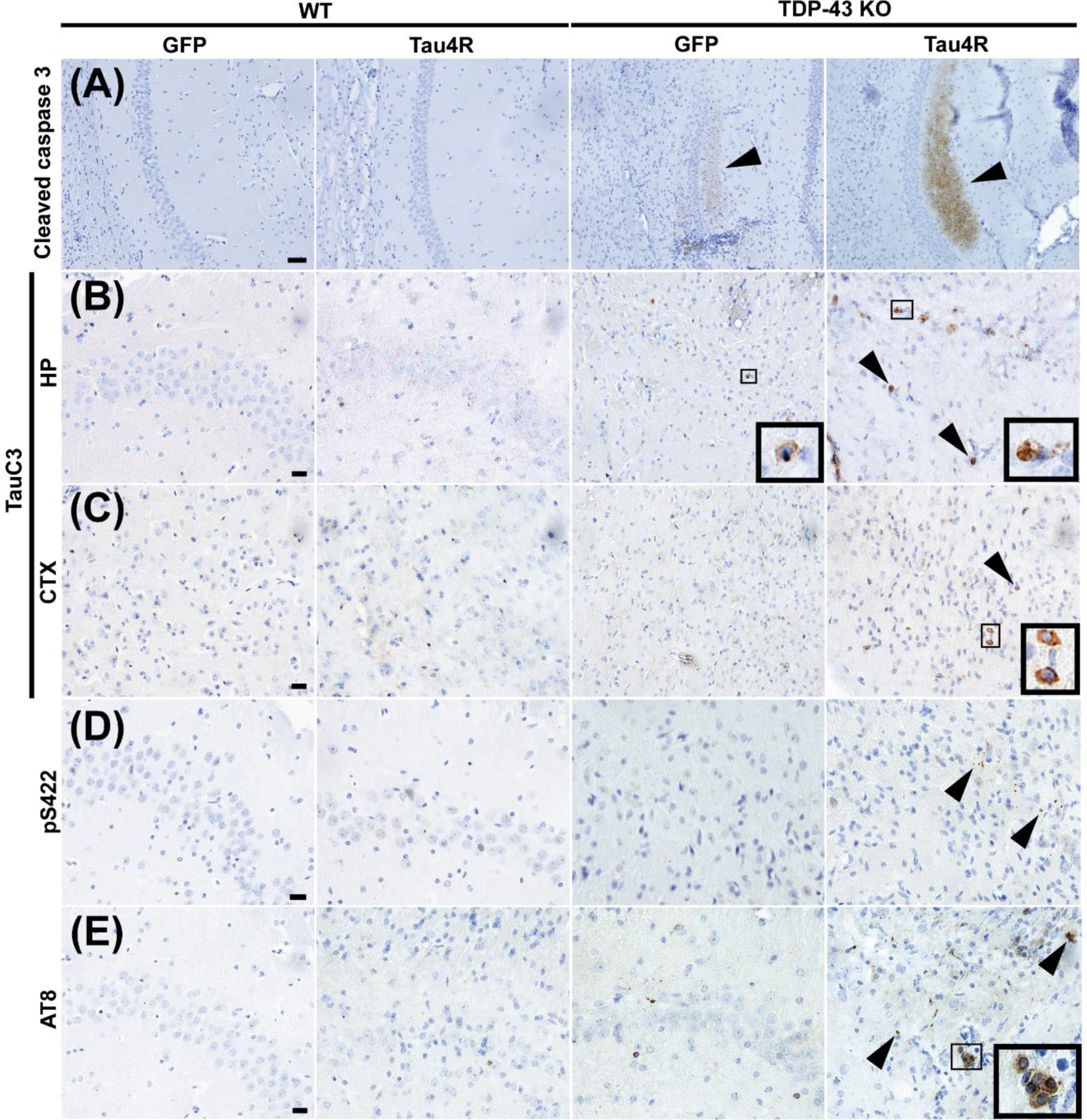
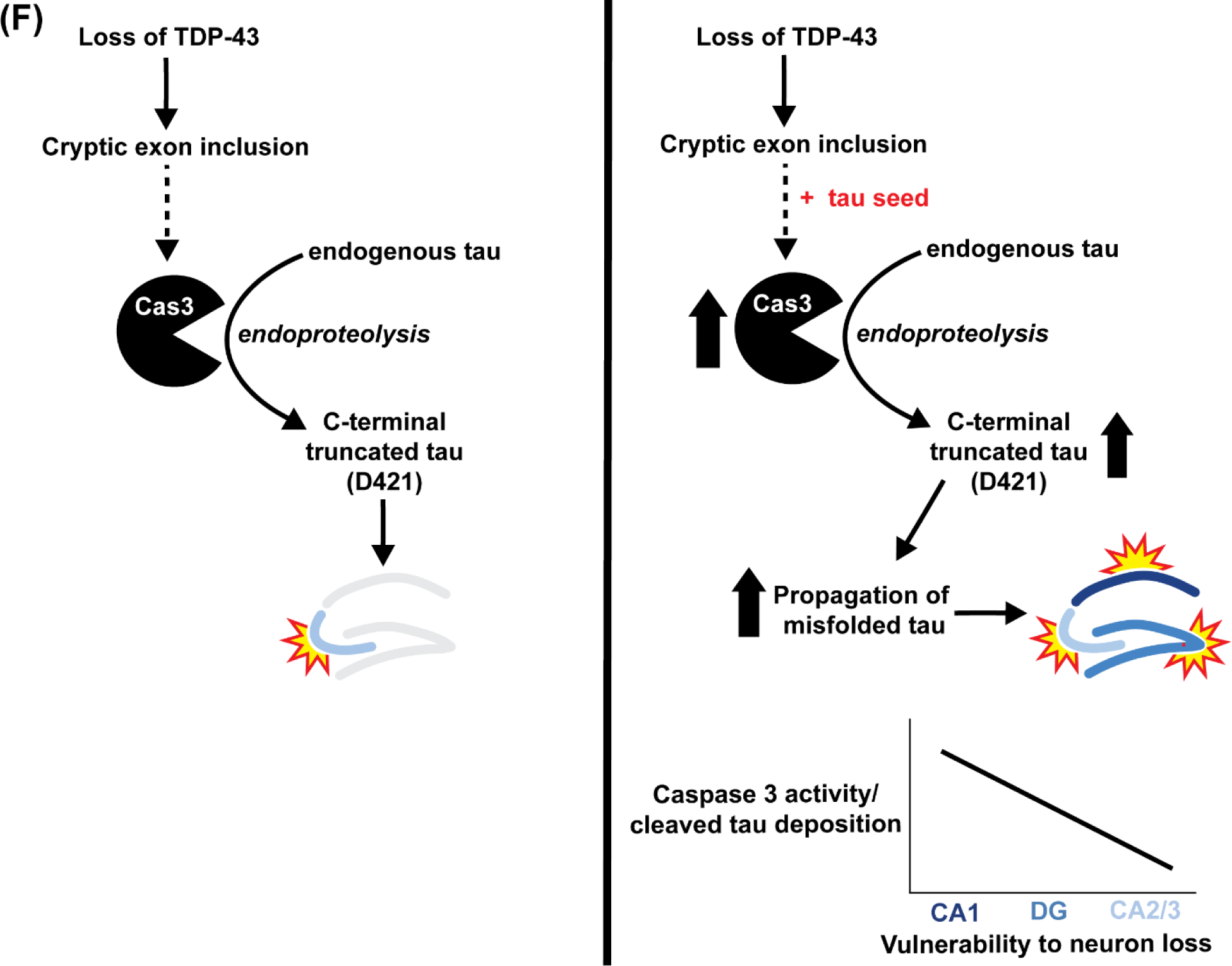
Caspase activation, tau cleavage and downstream tauopathy are further upregulated in mice lacking TDP-43 and expressing higher levels of hTauRD. **(A-E)** Immunohistochemistry of brain sections of 18-month-old WT mice injected with GFP (n=3), WT mice injected with Tau4R (n=4), TDP-43 KO mice injected with GFP (n=4), and TDP-43 KO mice injected with Tau4R (n=4) mice using antiserum against **(A)** cleaved caspase 3 in the CA1 region of the hippocampus (scale bar, 50µm), **(B-C)** caspase-cleaved tau (TauC3) in the CA2/3 region of the hippocampus and cortex directly above hippocampus (scale bar, 20 µm), and **(D)** phosphorylated S422 and **(E)** AT8 in the CA2/3 region (scale bar, 20μm). **(F)** Model depicting that loss of TDP-43 splicing repression of cryptic exons precedes caspase 3-mediated cleavage of endogenous tau and eventual degeneration of CA2/3 neurons upon loss of TDP-43 in the mouse hippocampus. In the presence of a tau seed, elevated caspase 3-mediated cleavage of tau drives pathological tau aggregation and extends neuronal vulnerability to dentate granule neurons and CA1 neurons.

### Intra-hippocampal delivery of a tau seed by AAV in *CaMKII-CreER;Tardbp^f^*^/^*^f^* mice elevates caspase 3-mediated tau cleavage and extends neuronal vulnerability to CA1

That the sensitivity of TDP-43 loss dependent increase of activated caspase 3 determines the vulnerability of neurons in CA3/2 or DG led us to speculate that CA1 neurons may require an even higher level of activated caspase to drive tauopathy. To test this notion, we unilaterally injected AAV-PhP.eB encoding hTauRD or GFP (Fig. 6A) to seed tauopathy in the dorsal hippocampus of 12-month-old *CaMKII-CreER;Tardbp^f^*^/*f*^ mice to drive higher accumulation of hTauRD and increase levels of activated caspase 3 (Fig. 6B). To validate viral infectivity, we employed a modified RNA *in situ* hybridization technique (BaseScope assay) using an RNA probe targeting an exon in the 5’-UTR of the AAV-PhP.eB vector. Viral RNA was detected in the hippocampus and the region of cortex above the hippocampus two months post injection (Fig. 6C). As a biochemical readout for *hTauRD* expression, we showed that *hTauRD* is only expressed in the right hippocampus and to a similar amount as endogenous tau (Fig. 6D). Additionally, we compared Tau4R expression between *Tau4R* mice and WT mice injected with AAV-Tau4R. Using an antibody against Tau4R, we demonstrate that AAV delivery results in higher levels of Tau4R expression in hippocampal neurons, including axons, compared to that in the Tau4R transgenic mice (Fig. 6E).

In agreement with our complementary transgenic *CaMKII-CreER;Tardbp^f^*^/*f*^*;hTau4R* model, *CaMKII-CreER;Tardbp^f^*^/*f*^ mice transduced with AAV-PhP.eB-hTauRD exhibited marked CA1, CA2/3, and DG neuron loss as well as hippocampal atrophy six months post injection (Figs. 6F-I). As expected, *CaMKII-CreER;Tardbp^f^*^/*f*^ mice injected with AAV-PHP.eB-GFP exhibited similar CA2/3 vulnerability due to loss of TDP-43 (Fig. 6H). Intriguingly, because of increased tau seeding through AAV delivery (Fig. 6E), neuronal loss in *CaMKII-CreER;Tardbp^f^*^/*f*^ mice injected with AAV-PhP.eB-hTauRD is evident in not only the CA2/3 and DG regions, as observed in *CaMKII-CreER;Tardbp^f^*^/*f*^*;hTau4R* mice, but also in the hippocampal CA1 region (Fig. 6G). These results suggest that increasing the level of activated caspase 3 in vulnerable neurons depleted of TDP-43 could drive tauopathy and CA1 cell death.

### Elevated caspase 3-dependent cleavage of endogenous tau accelerates tauopathy and death of neurons depleted of TDP-43 to include those in CA1

To determine whether increased caspase 3-mediated cleavage of tau underlies vulnerability of CA1 neurons to loss of TDP-43, we assessed caspase 3 activation in *CaMKII-CreER;Tardbp^f^*^/*f*^ mice injected with AAV-PhP.eB-hTauRD. As compared to AAV-PhP.eB-GFP injected mice as a control, we observed marked increase in activation of caspase 3 in *CaMKII-CreER;Tardbp^f^*^/*f*^ mice harboring AAV-PhP.eB-hTauRD (Fig. 7A). As expected, the degree of caspase activation correlated with caspase-cleaved tau observed in the hippocampus and cortex regions (Fig. 7B & C). Taken together, our data is consistent with the view that the vulnerability of neurons depleted of TDP-43 is dependent on the sensitivity of caspase 3 mediated cleavage of tau ranging from most sensitive ones in CA2/3, followed by those in the DG to the least sensitive in CA1. To test whether increased caspase 3-dependent cleavage of tau drives tauopathy and cell death in neurons within the CA1, we examined the extent of tauopathy in AAV-PhP.eB-hTauRD-treated *CaMKII-CreER;Tardbp^f^*^/*f*^ mice using antisera to pS422 and AT8. Six months post injection, we observed increased deposition of aggregated, hyperphosphorylated mouse (endogenous) tau in the hippocampus of *CaMKII*-*CreER;Tardbp^f^*^/*f*^ mice injected with AAV-PhP.eB-hTauRD as compared to that of controls (Fig. 7D & E). Our observations indicate that elevated levels of hTauRD promote greater activation of caspase 3-dependent cleavage of tau to facilitate pathological conversion of endogenous tau to drive neuronal death extending to those within CA1.

Together, our data support the model that while inclusion of TDP-43 dependent cryptic exons by itself triggers caspase 3 activation and CA2/3 neuron loss, TDP-43 depletion in the presence of a tau seed accelerates tauopathy in vulnerable neurons by promoting caspase 3-mediated endoproteolytic cleavage of tau, which in turn exacerbates the loss of increasingly vulnerable neurons in neurodegenerative disorders harboring co-pathologies of tau and TDP-43 (Fig. 7F). These mechanistic findings inform a novel therapeutic target/strategy to attenuate neuron loss occurring in MED with TDP-43 co-pathology, including AD-TDP.

## Methods

### Mouse Models

All the animal experiments were conducted as per the regulations of the Animal Care and Use Committee at Johns Hopkins University School of Medicine in accordance with the laws of the State of Maryland and the United States of America. We used the conditional knockout mouse model of TDP-43 by generating *CaMKIIα-Cre^ER^;Tardbp^f/f^* (TDP-43 KO) mice with loxP sites flanking *Tardbp* exon 3 (LaClair et al, 2016), creating a tamoxifen-induced recombination in excitatory forebrain neurons in adult mice. Through a two-stage breeding strategy (Sup Fig. 1), *CaMKIIα-Cre^ER^;Tardbp^f/f^*mice bred with *tTA;Tau4R* (Tau4R) line generated previously (Li et al 2016) and a cohort of *tTA;Tau4R;CaMKIIα-Cre^ER^;Tardbp^f/f^*(Tau4R;TDP-43 KO) mice on a C57BL/6J background, as well as littermate controls. Animals were genotyped at weaning, and mice were subsequently housed with one littermate of each genotype (5 mice/cage) when possible. Oral administration of tamoxifen citrate through diet was done in all animals (Harlan Teklad) at an average amount of 40 mg/kg/day for a 4-week period, beginning at 12 months-of-age. Studies in transgenic Tau4R;TDP-43 KO mice and littermate controls were carried out in brain tissue dissected out from 19-or 25-month-old mice.

For the AAV-PhP.eB-hTau4R/GFP injection model, we deleted TDP-43 at 11 months-of-age through the administration of tamoxifen feed. Mice were returned to normal feed as recovery for 2 weeks and then stereotaxic surgery was performed.

### Cell Culture and Transduction

The i3Neuron iPSC line was adapted from Michael Ward’s lab. iPSCs were cultured in Essential 8 media (Gibco, A1517001) on plates coated with Geltrex (Gibco, A1413301). iPSCs were differentiated into i3Neurons as described before (Tian et al., 2019). Lentivirus containing scrambled short hairpin RNA as non-targeting control (shScramble) or targeting TDP-43 (shTDP-43) as used before (Li et al., 2023), was transduced on day 5 after differentiation. Thereafter, neurons were harvested on day 8 to 16 or day 14 and 15 after differentiation, respectively, for cryptic exon analysis or immunoblot and immunostaining TDP-43, cleaved caspase 3 and cleaved tau.

### RT-PCR analysis

RNA was extracted from samples using the Monarch® Total RNA Miniprep Kit (New England Biolabs, T2010S) and cDNA was synthesized using the ProtoScript II First Strand cDNA Synthesis Kit (New England Biolabs, E6560L). Target sequences were amplified using a modified touchdown PCR protocol (Korbie and Mattick, 2008), primer pairs targeting wildtype with or without cryptic sequences (see table below) and Phusion Plus Green PCR Master Mix (Thermo Scientific, F632L). Upon cryptic exon inclusion, a larger band (CE) is expected. Amplified products were then separated by 1.5% agarose gel electrophoresis and visualized by ethidium bromide staining.

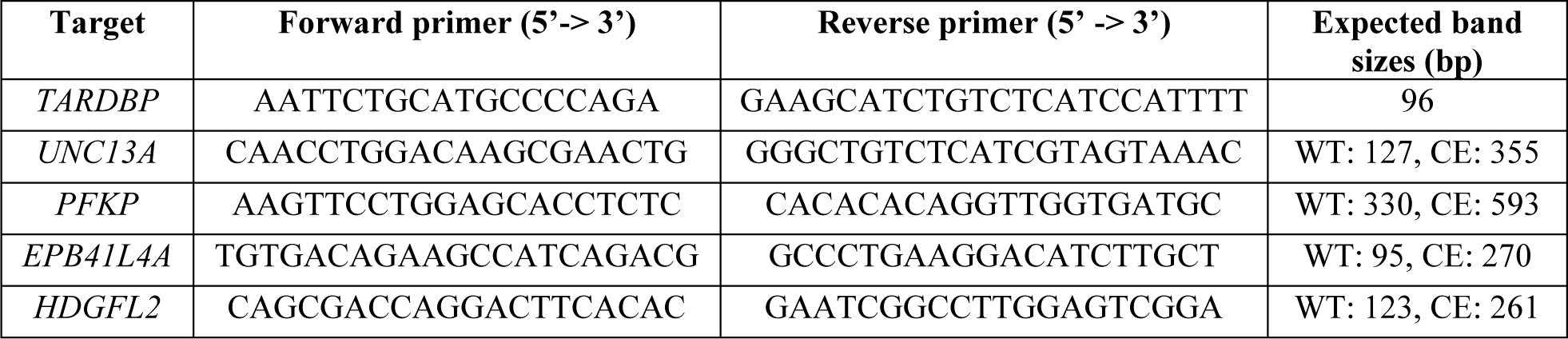

### Intraparenchymal stereotaxic injection

Mice were deeply anesthetized with isoflurane and fixed in a stereotaxic frame. The scalp skin of the animal was swabbed with povidone iodine and the skull was exposed via an incision. The skull was cleaned with hydrogen peroxide to allow for a craniotomy to be made above the right hippocampus using coordinates for dorsal hippocampus (2 mm frontal to lambda, 2 mm lateral to midline, and 1.5 mm depth) from the skull surface at the rate of 1µL/min as standardized before (Baghel et al., 2017). Unilateral injections were performed using a syringe (Hamilton) containing 5 µL of 2x10^13^ vg/mL AAV-PhP.eB-GFP or hTauRD. Upon completion, the needle was slowly withdrawn over 3 minutes. The skin was stapled and covered with 2% chlorohexidine to heal the incision.

### Immunoblotting

For protein blot analysis of cultured neurons, total protein was extracted with RIPA buffer (10 mM Tris-Cl (pH 8.0), 1 mM EDTA, 0.5 mM EGTA, 1% Triton X-100, 0.1% sodium deoxycholate, 0.1% SDS and 140 mM NaCl) containing 1X protease inhibitor cocktail (Roche, Indianapolis, IN). Samples were sonicated at 20 kHz for three five second pulses. Protein concentrations were determined by the BCA assay (Pierce Chemical Co., Rockford, IL) and equal amounts of protein lysates (∼20 μg per lane) were resolved on 4–12% Bis-Tris SDS–PAGE gels and then transferred to polyvinylidene difluoride membranes (Invitrogen, Carlsbad, CA). After blocking with 5% skimmed milk, the membranes were probed with the following antibodies: anti-TDP-43 N-terminus (1:2,000; 10782-2-AP; ProteinTech) and cleaved caspase 3 (1:1000; 5A1E; Cell Signaling Technology). Immunoblots were developed using enhanced chemiluminescence method (Millipore Corp., MA).

Total protein from the mouse hippocampus was extracted by homogenization in RIPA buffer. For mice injected with AAV, left and right hippocampi were dissected out separately and total protein was extracted. Protein concentrations were determined, and equal amounts of protein lysates (∼20 μg per lane) were resolved and transferred to polyvinylidene difluoride membranes. Then membranes were probed with the following antibodies: anti-human tau polyclonal antiserum KJ9A (1:5,000; A0024, Dako Cooperation, Carpinteria, CA) and rabbit anti-GAPDH antiserum (1:5,000; G9545, Sigma). Immunoblots were developed using enhanced chemiluminescence method (Millipore Corp., MA).

### Immunofluorescence Staining

For immunofluorescent staining of i3Neurons, neurons were fixed with an equal volume of 8% PFA to cell culture medium. Cells were then permeabilized with 0.1% Triton X. Blocking buffer (10% normal goat serum in PBS) was applied for an hour followed by primary antibodies overnight at 4°C. Secondary antibodies (Alexa Fluor 488, Alexa Fluor 594, Alexa Fluor 647) were added, and cells were stored in PBS for imaging.

The following primary antibodies were used: antiserum against TDP-43 C-terminus (1:1000; 12892-1-AP; ProteinTech), MAP2 (1:1000; 188004; Synaptic Systems), cleaved Caspase 3 (1:1000; D3E9; Cell Signaling Technology), Tuj1 (1:500; ab18207; Abcam), and cleaved tau (TauC3) (1:400; AHB0061, Invitrogen). The Zeiss Apotome Inverted Fluorescence Microscope (Zeiss, Germany) and the Zeiss LSM 800 AiryScan.2 were used for imaging.

To perform immunofluorescence staining on tissue, the paraffin brain sections were deparaffinized and antigen retrieval was performed using 10mM sodium citrate buffer to expose the epitope to the antibodies. Nonspecific binding of antibodies was eliminated by incubating with blocking buffer (1.5% normal goat serum in PBS with 0.1% Triton-X) for 1 hour. After blocking, the primary antibodies against Tau4R (E7T4F) (1:1000; 79327, Cell Signaling Technology); antiserum against cleaved caspase 3 were incubated overnight in humid chambers. Unbound antibodies were washed out and incubated with respective secondary conjugated fluorophores (Alexa Fluor 488 & Alexa Fluor 594). The Zeiss Apotome Inverted Fluorescence Microscope (Zeiss, Germany) was used for imaging.

### Histology and immunohistochemistry

The transgenic *Tau4R;TDP-43 KO* mice and their age matched littermates were sacrificed at 19 or 25 months of age for analysis. The injected mice were sacrificed 6 months post AAV-hTauRD injection. Mice were anesthetized with isoflurane and transcardially perfused with ice-cold PBS. Brains were dissected, postfixed in 4% PFA for 24 hours, embedded in paraffin and then sectioned with a microtome into 10µm thick sagittal sections.

For immunohistochemistry, the slides were deparaffinized and antigen retrieval was performed using 10mM sodium citrate buffer to efficiently expose the epitope to the antibodies. Nonspecific binding of antibodies was eliminated by incubating with blocking buffer (1.5% normal goat serum in PBS with 0.1% Triton-X) for 1 hour. Slides were then incubated in 0.3% H_2_O_2_ for 30 minutes to quench endogenous peroxidase. Primary antibodies were prepared in blocking buffer and applied overnight at RT followed by incubation with biotinylated secondary antibody for a 1 hour. Peroxidase labeled ABC reagent (Vector Laboratories) was applied for 30 minutes followed by signal development using 3,3 diaminobenzidine (DAB) (Vector Laboratories). Slides were counterstained with hematoxylin, dehydrated, cleared and mounted.

The following primary antibodies were used: antiserum against NeuN (1:1,000; MAB377, Merck); antiserum against TDP-43 N-terminus (1:1,000; 10782-2-AP, ProteinTech); phosphorylated S422 of tau (1:1,000; 44764G, Invitrogen, Carlsbad, CA); AT8 (1:1,000; MN1020, Invitrogen); antiserum against Tau4R (E7T4F) (1:1000; 79327, Cell Signaling Technology); antiserum against cleaved Caspase 3 (1:2000; D3E9; Cell Signaling Technology); antiserum against cleaved tau (TauC3) (1:200; AHB0061, Invitrogen).

The areas of hippocampus, cortex and cerebellum in sagittal sections at 2mm from the midline of the brains were measured using ImageJ. The numbers of healthy neurons in the CA1, CA2/3, and DG regions were counted manually using ImageJ.

### BaseScope-ISH assay

RNA in situ hybridization was performed using BaseScope Detection Reagent v2-RED Assay Kit (Cat#323900, Advanced Cell Diagnostics, Inc.), following manufacturer’s instructions. We used 3ZZ probes targeting an exon in the 5’-UTR of the AAV-PHP.eB vector (BA-CAG-promoter-O1-3zz-st-C1, Cat#1259091-C1, ACD Bio). A custom BaseScope probe (BA-Mm-Unc13a-O1-2EJ-C, Cat#1182491-C1, ACDBio) was used to detect the expression of transcripts harboring cryptic exon splice sites in Unc13a. To ensure RNA integrity, both positive (BA-Mm-Ppib, Cat#701071) and negative (BA-DapB, Cat#701011) control probes were used. Briefly, 10 µm tissue sections were deparaffinized and pre-treated with hydrogen peroxide, target retrieval buffer and protease IV and then hybridized with the target probes in a HybEZII oven (Advanced Cell Diagnostics, Inc.) for 2 hours at 40°C. The signals were amplified, and the sections were counterstained with hematoxylin. Images showing RNA puncta for each cryptic exon were captured using a Zeiss Apotome Inverted Brightfield Microscope (Zeiss, Germany).

### Statistical analysis

Graphs were generated using GraphPad Prism. Individual data points are also shown. One-way analysis of variance (one-way ANOVA) for multiple comparisons was performed using the GraphPad Prism software (La Jolla, CA, USA). In all tests, values of p<0.05 were considered significant. The number of biological replicates and details of statistical analyses are provided in the figure legends.

## Discussion

Alzheimer’s Disease (AD) and AD-Related Dementias (ADRD) are a group of progressive and complex neurodegenerative disorders with mid-to late-life onset, including Lewy body dementia (LBD), frontotemporal dementia (FTD), limbic-predominant age-related TDP-43 encephalopathy (LATE), or multiple etiology dementia (MED) such as AD with co-pathology of a-synuclein and/or TDP-43 (Josephs et al., 2014; Josephs et al., 2015; Robinson et al., 2018; Neumann et al., 2006; Probst al., 1996; Stefanis, 2012; Nelson et al., 2019). To clarify disease mechanisms, identify therapeutic targets, and validate therapeutic strategies for these complex dementias, a critical unmet need is the availability of appropriate model systems that mimic key aspects of these co-pathologies, as they account for upwards of 75% of all cases of ADRD. The acuity of this problem was exposed in recent FDA-approved drugs targeting only β-amyloidosis (Drug Approval Package: Aduhelm (aducanumab-avwa) (fda.gov)) as many participants failed to respond positively for a drug that does not address the pathophysiology of tau, α-synuclein or TDP-43.

The presence of these co-pathologies is thought to drive accelerated neurodegeneration and steeper cognitive deficit, which raises an important question as to how these co-pathologies contribute to disease pathogenesis. Since TDP-43 co-pathology has not only been observed with tauopathy in AD brains, but also in cases of FTLD (Kim et al., 2018), CBD (Koga et al., 2018), PART (Josephs et al., 2017) and PSP (Yokota et al., 2010), clarifying how this RNA binding protein influences tauopathy would provide insight into pathogenic mechanisms and novel therapeutic targets. We previously discovered that a major function of TDP-43 is the transcriptome-wide repression of non-conserved cryptic exons, which is impaired in neurodegenerative diseases exhibiting TDP-43 pathology (Ling et al., 2015, Donde et al., 2019). That loss of TDP-43 function reflects TDP-43 pathology and drives neuron loss is strengthened through the identification of disease-associated TDP-43 dependent cryptic exon targets (Ma et al., 2022, Brown et al., 2022, Klim et al., 2019; Melamed et al., 2019), nuclear clearance without TDP-43 cytoplasmic aggregates in AD cases with TDP-43 pathology (Sun et al., 2017), and the discovery that loss of TDP-43 splicing repression occurs early in disease, including the pre-symptomatic stage (Irwin et al., 2024; Chang et al., 2023; Seddighi et al., 2024). These studies support the view that TDP-43 loss of function during the presymptomatic stage of disease may influence tauopathy to accelerate neuron loss in MEDs exhibiting the co-pathologies of tau and TDP-43. By crossing mice lacking TDP-43 in forebrain neurons to a tau-seeding model, we generated a novel mouse model of MED designed to clarify the influence of TDP-43 pathology on exacerbated neuron loss in human tauopathies with co-pathology of TDP-43.

Our findings are consistent with a model in which inclusion of TDP-43 dependent cryptic exons in forebrain neurons exacerbates tauopathy-dependent atrophy of the hippocampus by sensitizing vulnerable neurons to caspase 3-dependent endoproteolysis of tau: from most vulnerable pyramidal neurons in CA2/3 to granule neurons in the dentate and to the least in CA1 (Fig. 7F). This view is supported by our two complementary approaches to increase levels of caspase 3-cleaved tau by modestly or robustly seeding tau, respectively, using our Tau4R transgenic mice (Figs. 3-5) or AAV-PHP.eB-TauRD (Figs. 6 & 7); in the former, higher levels of caspase 3-cleaved tau achieved by *CamKII*-*CreER;Tardbp^f^*^/*f*^*;hTau4R* as compared to *CamKII*-*CreER;Tardbp^f^*^/*f*^ mice led to neuron loss not only within CA2/3 but also include those in the dentate; whereas the latter accumulated greatest amount of caspase 3-cleaved tau in the hippocampus of AAV-TauRD injected *CamKII*-*CreER;Tardbp^f^*^/*f*^ mice to drive neuron loss to include even the least vulnerable ones found in CA1. These findings establish that levels of caspase 3-cleaved tau determine the selective vulnerability of neuron lacking TDP-43 that is sensitive to levels of caspase 3-cleaved tau in the mouse hippocampus. Coupled with our observation that inclusion of cryptic exons preceding caspase 3-mediated endoproteolysis of tau is found in human iPSC-derived cortical neurons lacking TDP-43 (Fig. 2), our discoveries suggest that loss of TDP-43 promotes endoproteolysis of endogenous tau by caspase to exacerbate tauopathy-and age-dependent neuron loss is the pathogenic mechanism underlying the exacerbated neurodegeneration observed in human tauopathies with co-pathology of TDP-43. Importantly, we also show in our mouse of MED that loss of TDP-43 leads to incorporation of cryptic exons and subsequent activation of caspase 3 (Fig. 1). These observations suggest that TDP-43 splicing repression is a novel therapeutic target to attenuate neurodegeneration in MED with co-pathologies of tau and TDP-43, a therapeutic strategy which can be validated in our novel mouse model of MED. We recently validated such an AAV gene therapy strategy to complement the loss of TDP-43 splicing repression (Ling et al., 2015; Donde et al., 2019). Since loss of TDP-43 splicing repression function is upstream of caspase 3-cleaved tau, it is likely that such therapeutic approach could provide clinical benefit for patients harboring these two co-pathologies. Testing this AAV gene therapy approach or in combination with a tau drug, such as an ASO against tau (currently in Phase II clinical trials), using our novel *CaMKII-CreER;Tardbp^f^*^/*f*^*;hTau4R* mouse and human models of MED will be instructive.

Based on our proposed model (Fig. 7F), there are several strong predictions that would further establish the pathogenic mechanism whereby loss of TDP-43 function exacerbates neurodegeneration in MED with co-pathology of tau and TDP-43. First, our studies have not directly considered the role of β-amyloidosis in relationship to loss of TDP-43 function that occurs in cases of AD-TDP. We and others previously found that the β-amyloid plaque is necessary for the pathological conversion of tau (Li et al, 2016; He et al., 2018; Balusu et al, 2023) and identified disease stage-specific activation of microglia by either β-amyloid plaque or tau aggregates (Keren-Shaul et al., 2017; Masuda et al., 2019; Chen et al, 2020; Kim et al, 2022). That the amyloid-β plaque in the presence of TDP-43 depletion also promotes caspase 3 activation (LaClair et al, 2016) would suggest the idea that β-amyloidosis might facilitate and exacerbate tauopathy in neurons lacking TDP-43 through this same mechanism, a prediction that can be addressed using mouse models of β-amyloidosis (Li et al., 2016). Future studies will be necessary to delineate whether and how β-amyloid plaques and TDP-43 loss of function may converge on the caspase 3-dependent cleavage of tau to drive tauopathy.

Another strong prediction is that the inhibition of caspase 3 activation by loss of TDP-43 function using pharmacological or genetic approach would attenuate tauopathy in either of our mouse or human models of MED. Supporting this view are previous studies documenting: 1) inhibition of endoproteolysis tau (at D421) using the specific antibody TauC3 (which recognizes the truncated tau neoepitope) substantially prevents the seeding competency of AD brain lysate with high molecular weight tau aggregation in cultured cells (Nicholls et al, 2017); 2) overexpression of caspase 3 in neuronal cells increases tau phosphorylation and tauopathy (Chu et al., 2017; Biundo et al., 2017); 3) co-expression of humanized caspase 6 and truncated tau (TauΔD421) fails to lead to tauopathy or neurodegeneration in the aged mouse brain (Noël et al., 2022); 4) mice expressing C-terminal truncated tau at D421 exhibit synaptic and cognitive impairments through oligomerization of tau (Kim et al., 2016); 5) AAV delivery of C-terminal truncated tau to mouse brains results in hyperphosphorylation and oligomerization of tau leading to aggregated tau tangles (Loon et al., 2022); and 6) caspase inhibition attenuates (Theofilas et al., 2023) caspase 6 cleaved tau in iPSC neurons derived from a *MAPT-*V337M associated FTD patient.

Based on our previous finding that the central role of TDP-43 is to repress the splicing of cryptic exons (Ling et al, 2015; Donde et al, 2019), and inclusion of cryptic exons occurs before caspase 3 activation, we hypothesize that, due to loss of TDP-43 function, cryptic exon inclusion can promote caspase 3-mediated endoproteolysis of tau to drive tauopathy. We speculate that inclusion of multiple cryptic exons, as opposed to any specific ones, underlies activation of caspase 3. From first principle, cryptic exons are nonconserved (Ling et al., 2015; Jeong et al., 2017), yet caspase 3 cleaved tau occurred in both mouse and human models lacking TDP-43. TDP-43 associated cryptic exons are linked to disease and elevated caspase 3-mediated tau cleavage is also observed in both systems would suggest a common molecular mechanism. One overlapping TDP-43 target between mice (Jeong et al., 2017) and humans is *UNC13A* (Seddighi et al., 2023; Melamed et al., 2019; Klim et al., 2019; Brown et al., 2022; Ma et al., 2022), where cryptic exon inclusion results in nonsense-mediated decay of *UNC13A* transcripts and reduced protein levels. These observations suggest that *UNC13A* alone could be a potential target to reverse the caspase 3 activation-mediated events. In contrast, the caspase-mediated event could also be the consequence from the combinatorial effect of a distinct set of cryptic exons in neurons lacking TDP-43. Future studies will be necessary to resolve these two possibilities. Nevertheless, our discoveries establish a novel mouse model of MED exhibiting the co-pathology of tau and TDP-43, disclose a pathogenic mechanism underlying exacerbated neuron loss and offer a potential therapeutic strategy designed to either complement the loss of TDP-43 function (Ling et al., 2015; Donde et al., 2019) and/or a combination therapy to target cryptic exon splicing and tau (such as ASO against tau) in vulnerable neurons of these devastating human disorders currently without any effective therapy.

## Acknowledgments

We thank T. Melnikova and B. Kim for technical support.

## Authors Contributions

M.S.B. and G.B. conceived, designed and coordinated the studies, performed the histological, biochemical, and statistical analysis and wrote the manuscript. M.T., T.C., X.K.C., and I.D.R. performed immunohistochemical analysis of Tau4R;TDP-43 KO and TDP-43 injected with AAV-PhP.eB-GFP/Tau4R mice. A.P.M. performed BaseScope analysis of *Unc13A* and AAV-Tau4R in TDP-43 KO and injected mice. G.B., A.L.C. and Y.Y. performed and analyzed the TDP-43 KD studies in human iPSCs derived neuron. S.R.M. has performed brain sectioning and helped with neuron quantification. P.C.W. conceived and designed studies and edited the manuscript. S.S. and T.L. edited the manuscript, and all authors read and approved the final manuscript.

## Author Statement

This article was prepared while Dr. Tong Li was employed at The Johns Hopkins University School of Medicine. The opinions expressed in this article are the author’s own and do not reflect the view of the National Institute on Aging, the National Institutes of Health, the Department of Health and Human Services, or the United States government.

---

**Supplementary Figure 1:**
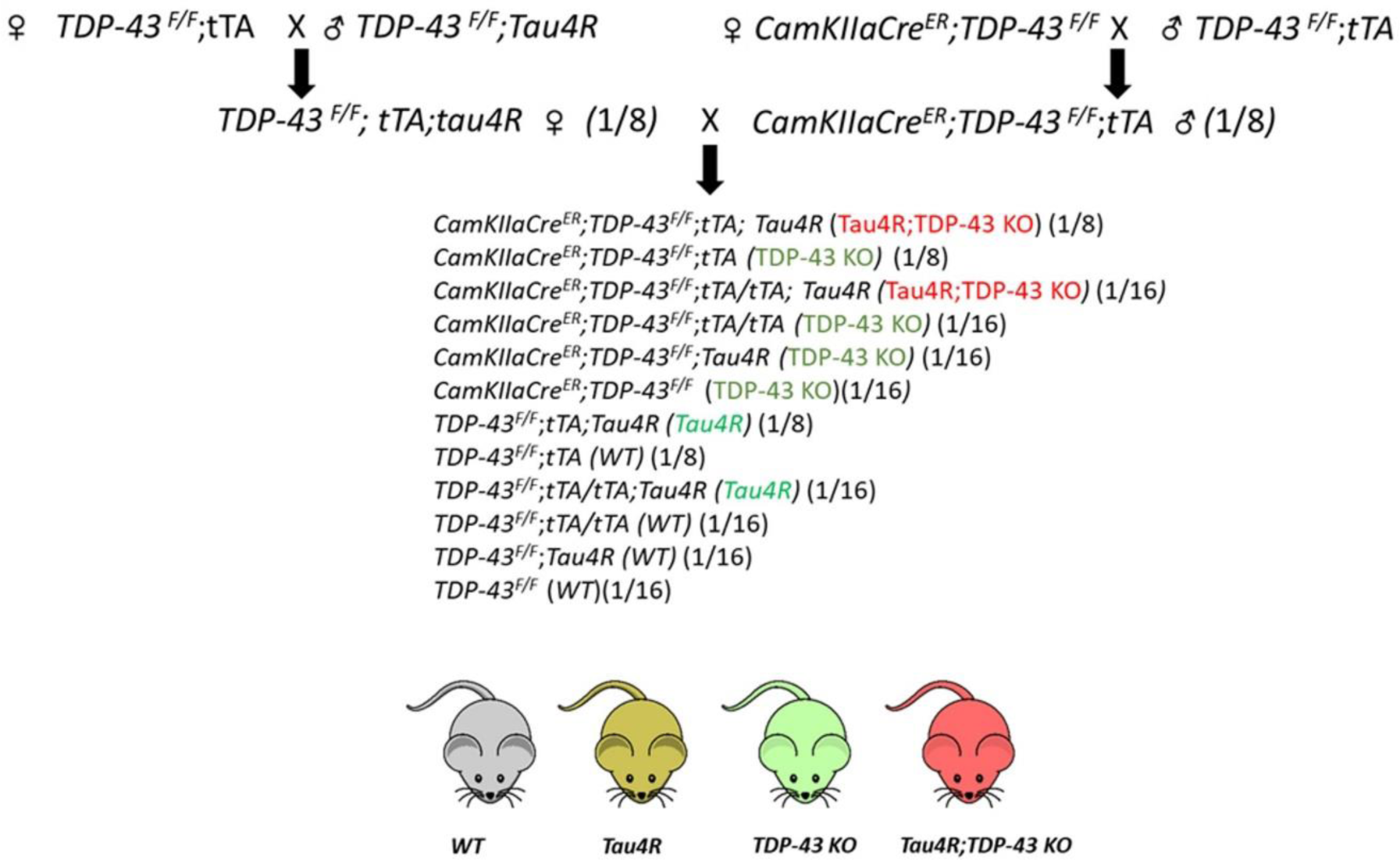
Breeding strategy for generating *CaMKII-CreER;Tardbp^f/f^*;*hTau4R* mice and littermate controls. (#) Indicates the Mendelian frequency of pups from each litter expected to carry the genotype of interest

**Supplementary Figure 2:**
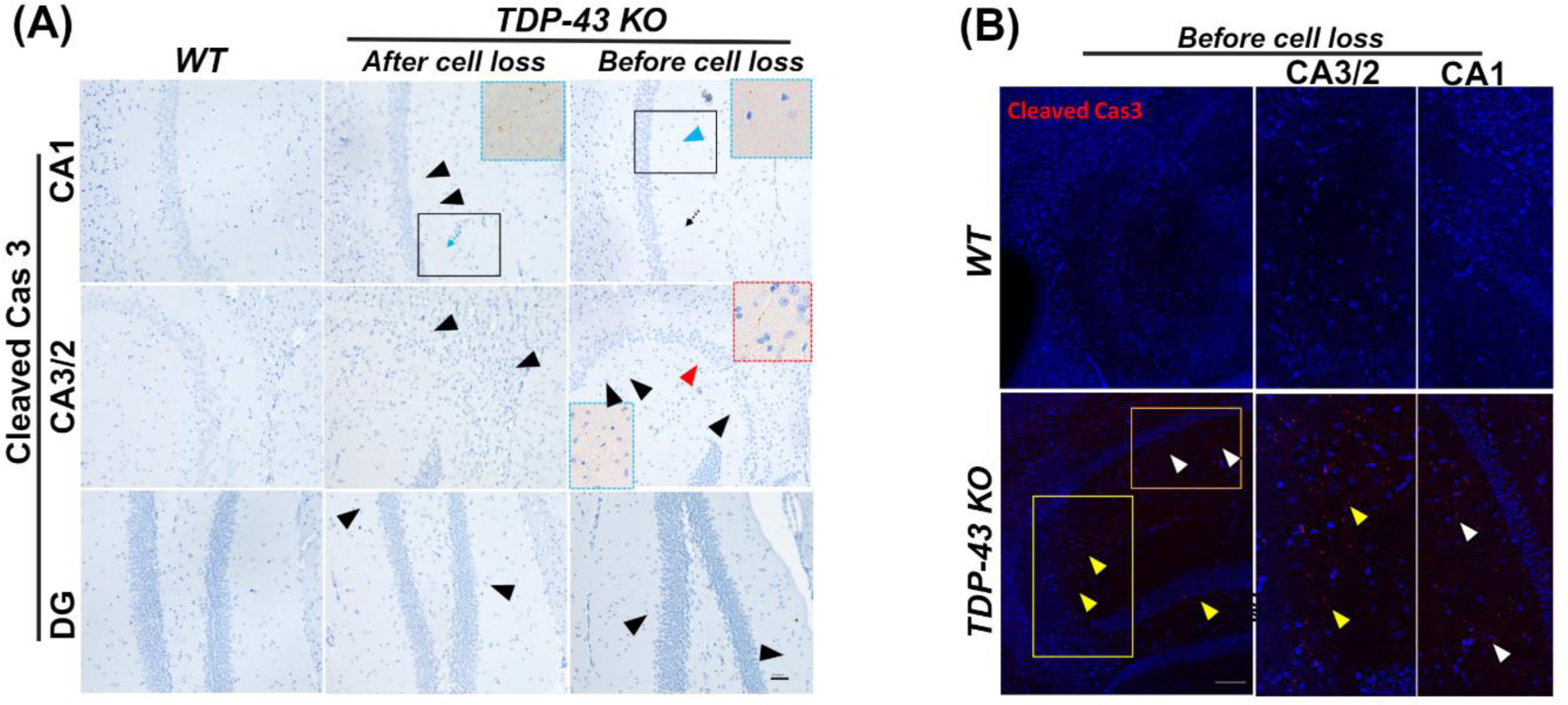
(A) Immunohistochemical analysis of. *WT* (left column) and *TDP-43 KO* mice (middle column; after cell loss) and before cell loss (*TDP-43 KO* 4 M, right column) using antiserum against cleaved caspase 3 in the CA1, CA3/2 and DG subregion of the hippocampus, arrow heads indicating the immunoreactivity enlarged view in inset (scale bar, 50μm). **(B)** Immunofluorescence of cleaved caspase 3 corroborating the immunohistochemistry before the neuron loss 2M after deletion (Figure 1 B), showing cleaved caspase 3 immunofluorescence in CA1, CA3/2 and DG subregions as arrow heads indicate (left column) enlarged view in middle (CA3/2) and right column (DG) (scale bar, 100μm).

**Supplementary Figure 3:**
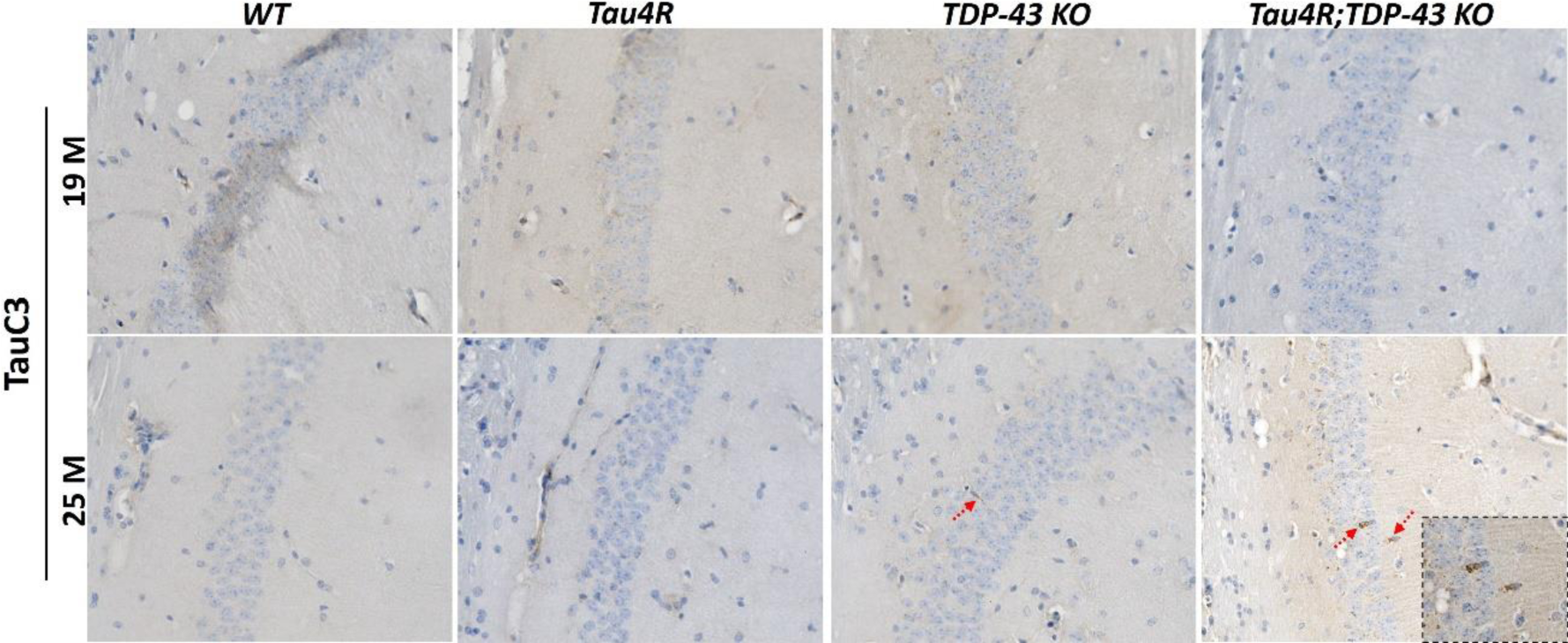
Immunohistochemical analysis of cleaved tau (TauC3) in brain sections of 19 and 25 M old *WT* (n=3), *Tau4R* (n=4), *TDP-43 KO* (n=5) and *Tau4R;TDP-43 KO* mice (n=4) in the CA1 region of the hippocampus using antiserum against truncated tau. The arrowheads show the cleaved tau signal in *TDP-43 KO* and *Tau4R;TDP-43 KO* aged (25 M), but not in WT and Tau4R transgenic mice.

**Supplementary Figure 4:**
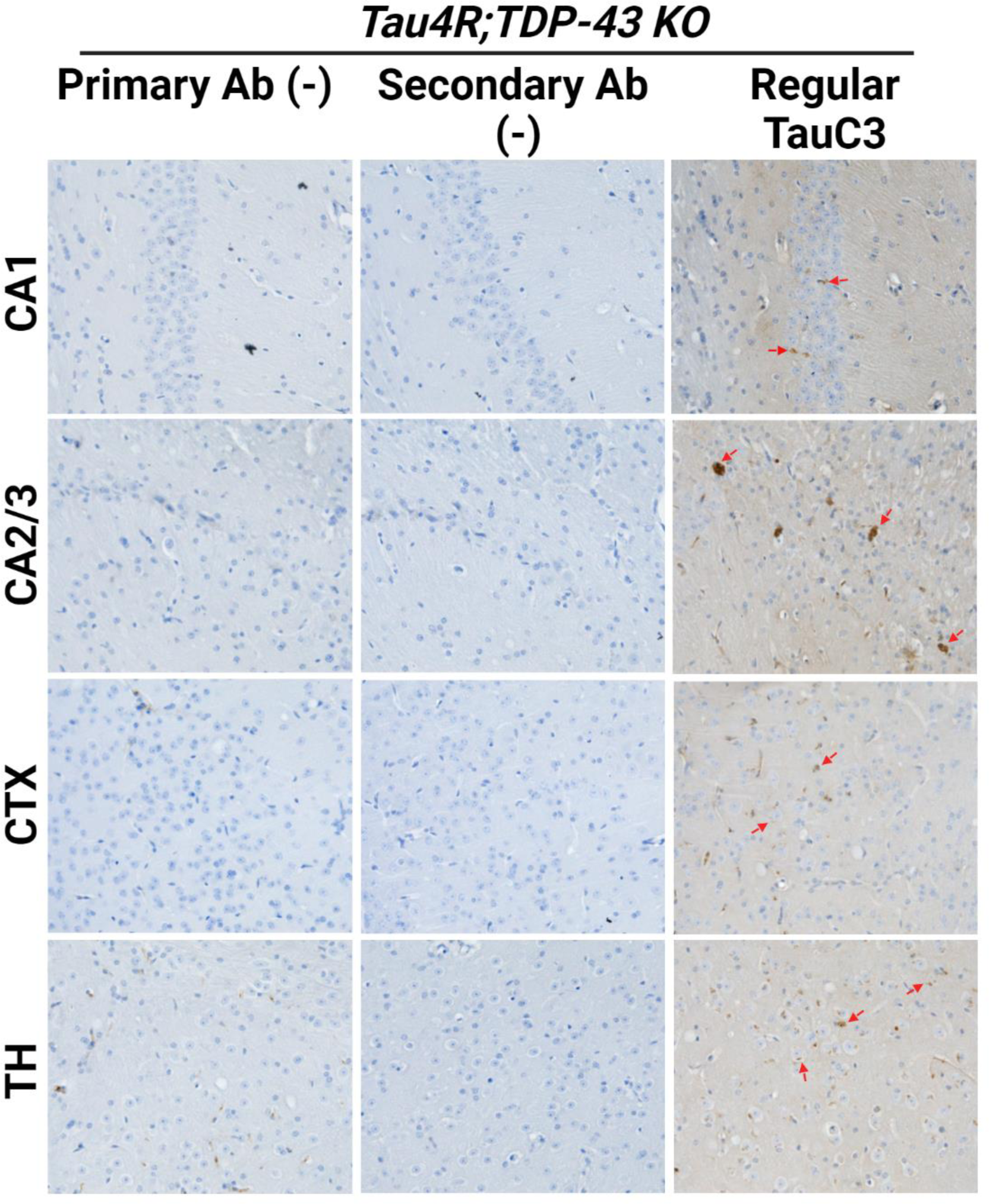
Testing the specificity of TauC3 immunoreactivity: Immunohistochemical analysis of cleaved tau (TauC3) in brain sections of 25 M old *Tau4R;TDP-43 KO* mice; shows immunoreactivity in regular staining with TauC3 antibody in different brain regions (Right column). The arrowheads show the cleaved tau signal. The without primary antibody (Left column) and secondary antibody (Middle column) incubation show no immunoreactivity (brown staining).

**Supplementary Figure 5:**
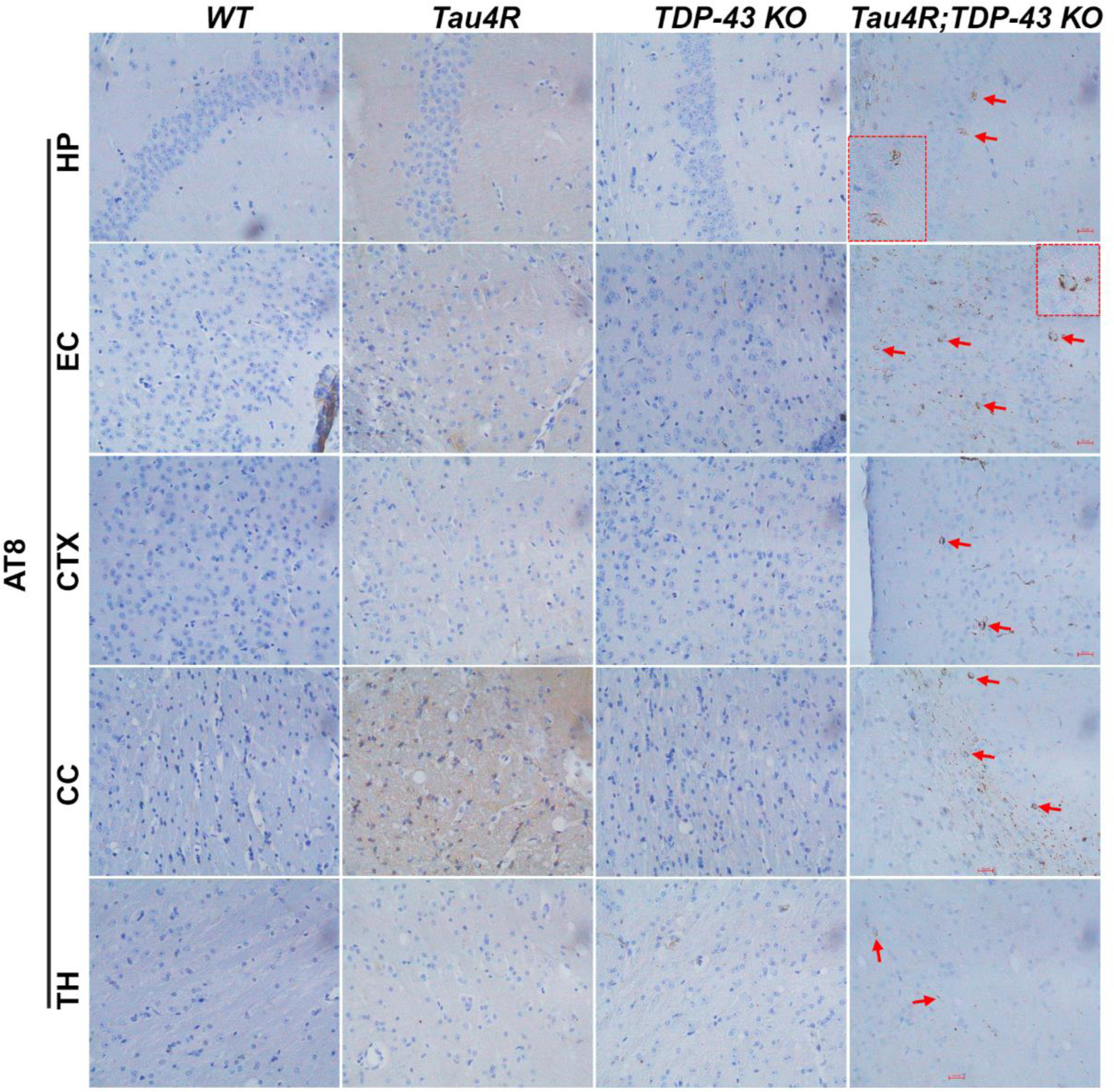
Immunohistochemistry of pathological tau (AT8) in brain regions of *Tau4R;TDP-43 KO* old mice (25 M) mice, pathological tau tangles could be detected. The brain regions are entorhinal cortex (EC), cerebral cortex (CTX), hippocampus (HP); carpus callosum (CC) and thalamus (TH). Scale bar, 20μm.

**Supplementary Figure 6:**
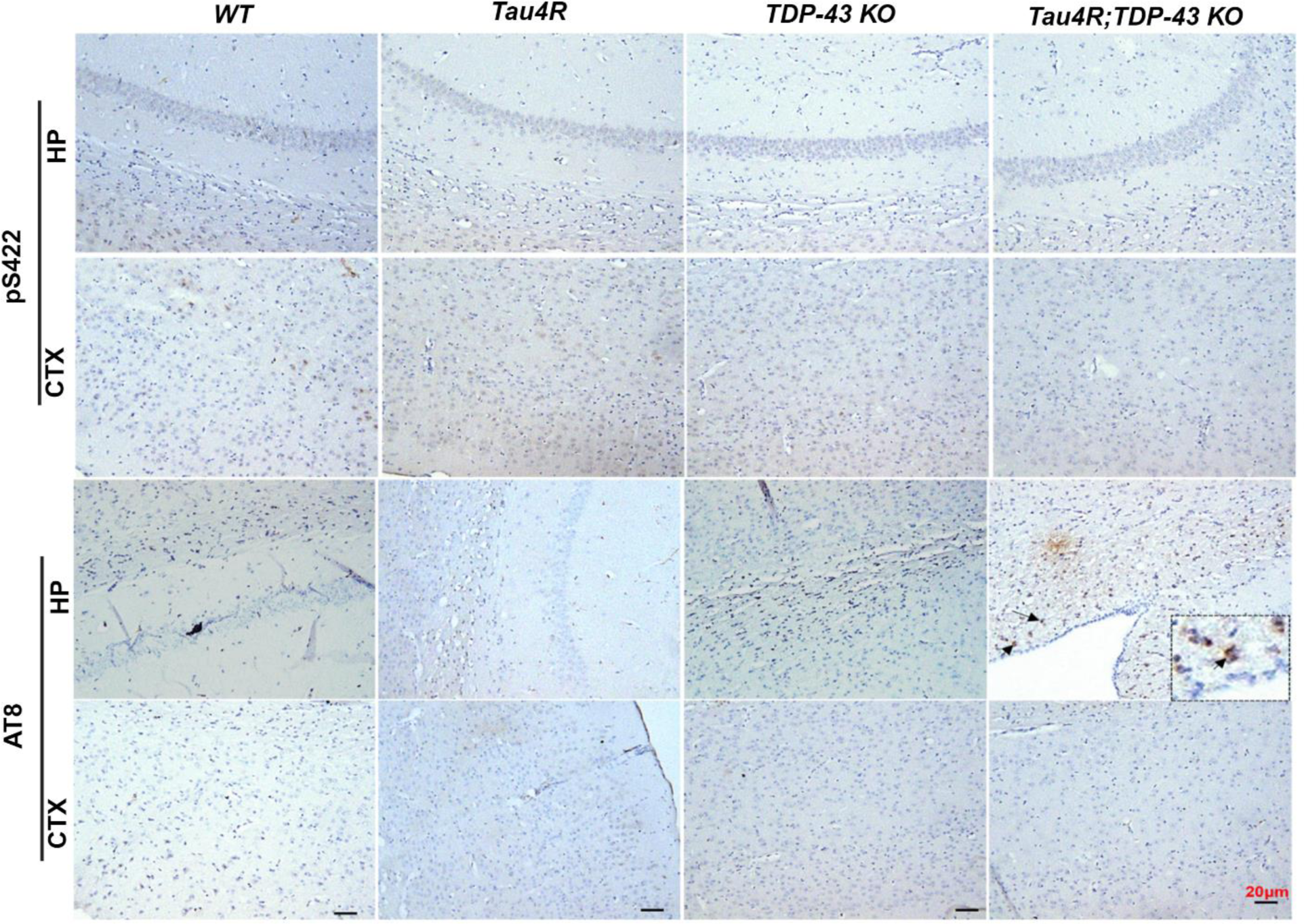
Immunohistochemistry analysis in hippocampus and cortex using antibodies specific to endogenous phosphorylated tau, pS422 (upper panel) and AT8 (lower panel). Arrowhead shows probably some brown staining of AT8 but not pS422 around the ventricle side of hippocampus in *Tau4R;TDP-43 KO* mice (n=5) at the age of 19 M (scale bar, 20μm), but staining does not look like pathological tau. The sections were counterstained with hematoxylin (blue).

**Supplementary Figure 7:**
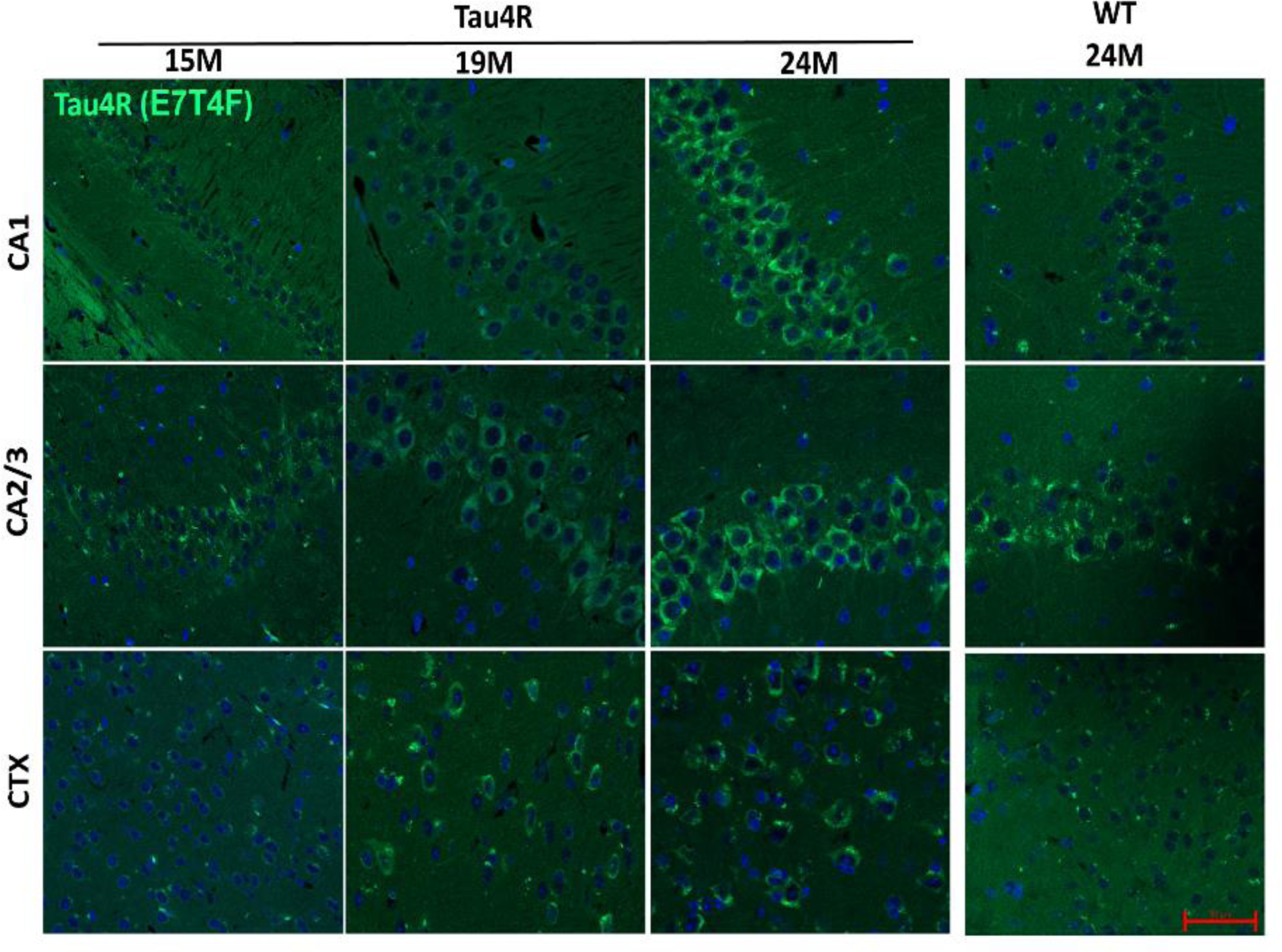
Immunofluorescence analysis corroborated the immunohistochemistry of tau4R fragment and showed an age-dependent increase in the accumulation of hTau4R fragment using 4R repeat recognizing antibody (Tau4R;E7T4F) in subregions of HP and CTX of *Tau4R* mice (aging 15, 19 & 24 M), but not in *WT* mice right panel (24M) (Scale bar, 50 μm)

